# Chromatin organization controls nuclear stiffness

**DOI:** 10.1101/2025.03.14.643219

**Authors:** Hector Romero, Anahid Amiri, Maruthi K. Pabba, Hui Zhang, Veronika Berg, Maria Arroyo, Paulina Prorok, Nina Trautwein, Bodo Laube, Christian Dietz, Robert W. Stark, M. Cristina Cardoso

## Abstract

Cellular differentiation is driven by epigenetic modifiers and readers, including the methyl CpG binding protein 2 (MeCP2), whose level and mutations cause the neurological disorder Rett syndrome. During differentiation, most of the genome gets densely packed into heterochromatin, whose function has been simplistically viewed as gene silencing. However, gene expression changes reported in mutations leading to Rett syndrome have failed to be a predictor of disease severity. Here, we show that MeCP2 increases nuclear stiffness in a concentration dependent manner and dependent on its ability to cluster heterochromatin during differentiation. MeCP2-dependent stiffness increase could not be explained by changes in the expression of mechanobiology-related genes, but we found it is disrupted by Rett syndrome mutations and correlated with disease severity. Our results highlight the impact of chromatin organization in the mechanical properties of the cell as an alternative or complementary mechanism to changes in cytoskeleton components.

**Graphical abstract:** 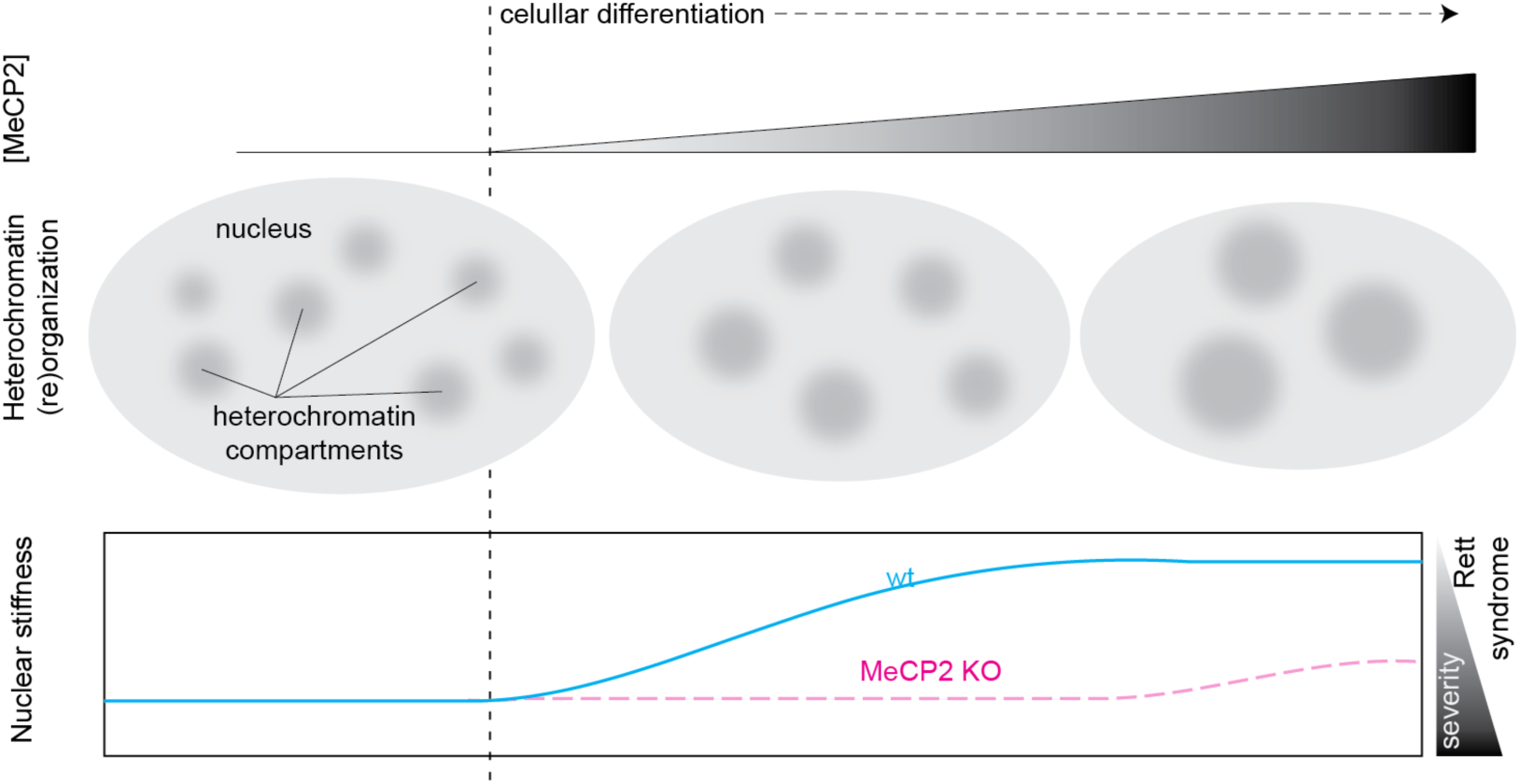

## Introduction

The (micro)environment surrounding cells is fundamental to understanding the function of these cells. In recent years, the importance of the physical properties of the environment, which drive different biological processes, including proliferation(*1*), mobility(*2*), and differentiation(*3*–*8*) is gaining special relevance. Changes in the physical properties of the tissues have been also related to the effects of aging (*9*) and disease (*9*, *10*).

Physically mediated processes are very complex to study in biological samples. Not only are there differences in the stiffness between tissues (*11*), but also among regions in the same tissue, such as different areas of the brain (*12*). Indeed, even different types of cells located in the same environment respond differently to the same stimulus, which suggests the presence of mechanisms that regulate responses to physical signals. Most studies try to answer this question through the investigation of the changes in cytoskeleton and/or membrane components (*11*–*18*). Interestingly, although the nucleus has been considered to play a major role in mechanical responses (*2*, *17*, *19*–*22*), the contribution of the chromatin organization to the differential responses to mechanical stress has been rarely addressed.

Mostly, the nuclear stiffness has been characterized to be dependent on the laminA concentration (*11*) and its contribution related to the linker of nucleoskeleton and cytoskeleton (LINC) complex (*23*). However, studies showed that substrate stiffness (*24*–*26*), topographical cues (*27*), and cellular geometry (*28*–*30*), modulate nuclear organization, chromatin remodeling and gene expression. These findings indicate that mechanical cues are diverse in nature and selectively regulate the epigenome, with significant implications for physiology and therapeutic applications.

Each human cell contains meters of DNA packed into an ovoid nucleus with 5-20 µm diameter (*20*). To make such a dense packing possible, DNA is condensed into chromatin. However, the condensation of the chromatin is not homogenous, leading to membraneless compartments within the nucleus: euchromatin (more open and accessible chromatin) and heterochromatin (condensed chromatin). These compartments are highly dynamic and undergo significant reorganization during cell differentiation (*31*–*34*). This reorganization is directed by epigenetic marks and driven by epigenetic readers. One prominent reader is MeCP2, whose levels are dependent on cell differentiation (*33*, *34*). MeCP2 induces the clustering of heterochromatin in a concentration-dependent manner (*35*), and changes in its levels or mutations in the MeCP2 gene are linked to diseases, with special relevance of the neurological disorder Rett syndrome (OMIM: #312,750) (*36*–*38*), as MeCP2 is especially abundant in neurons (*39*, *40*).

Taking together the importance of the nucleus in the cell response to mechanical stress and the possibility of changing the organization of the chromatin within the nucleus, we hypothesize that nuclear stiffness is regulated not only by the cytoskeleton but also through dynamic chromatin organization, being dysregulated in disease. In this study, we investigated the effect of this epigenetic reader and chromatin organizer (MeCP2) on nuclear stiffness using atomic force microscopy (AFM) to measure the elastic modulus of nuclei and cells, alongside analyzing chromatin organization. We aimed to determine how MeCP2 level and chromatin clustering influence nuclear stiffness, particularly in the context of cellular differentiation and disease models, including Rett syndrome, and to assess the role of mechanotransduction pathways in these processes. Our results suggest that MeCP2 is a major factor increasing nuclear stiffness in neuronal differentiation systems, and that this function is relevant for the severity of the Rett syndrome derived from MeCP2 mutations.

## Results

### The stiffness of the nucleus is independent of the cytosolic cytoskeletal components

We first purified the nuclei to remove all cytosol components, including the major cytoskeleton components. Since AFM involves direct interactions between the cantilever and the sample, we found that the nuclei exhibited drift during image acquisition. To address this issue, we embedded the nuclei in a soft agarose gel, which stabilized the sample and prevented movement. This approach allowed us to accurately measure the elastic modulus of the purified nuclei. All measurements in this study were conducted under identical conditions, consistent sample preparation, experimental parameters (tip speed, setpoint, resolution), and methodology, ensuring comparability across samples.

We compared the elastic modulus measurements obtained from whole cells and specifically in the nuclear region to those measured in purified nuclei isolated from the same cell type (Figure 1, Table 1). Our results showed that: i) the nuclear region within the intact cells exhibited higher elastic modulus values, with a median value of 1.5 kPa, compared to that of the whole cell, with a median value of 0.9 kPa; and ii) the purified nuclei displayed elastic modulus values comparable to those of nuclear region in intact cells, with a median value of 1.4 kPa. The similarity in stiffness distributions between purified nuclei and the nuclear region of intact cells suggests that potential dehydration effects during nuclei purification, which could influence elastic modulus measurements (*22*, *41*) are minimal under our experimental conditions. However, in the absence of cytosolic cytoskeletal components, the distribution of elastic modulus values in purified nuclei was more homogeneous.

**Figure 1.**
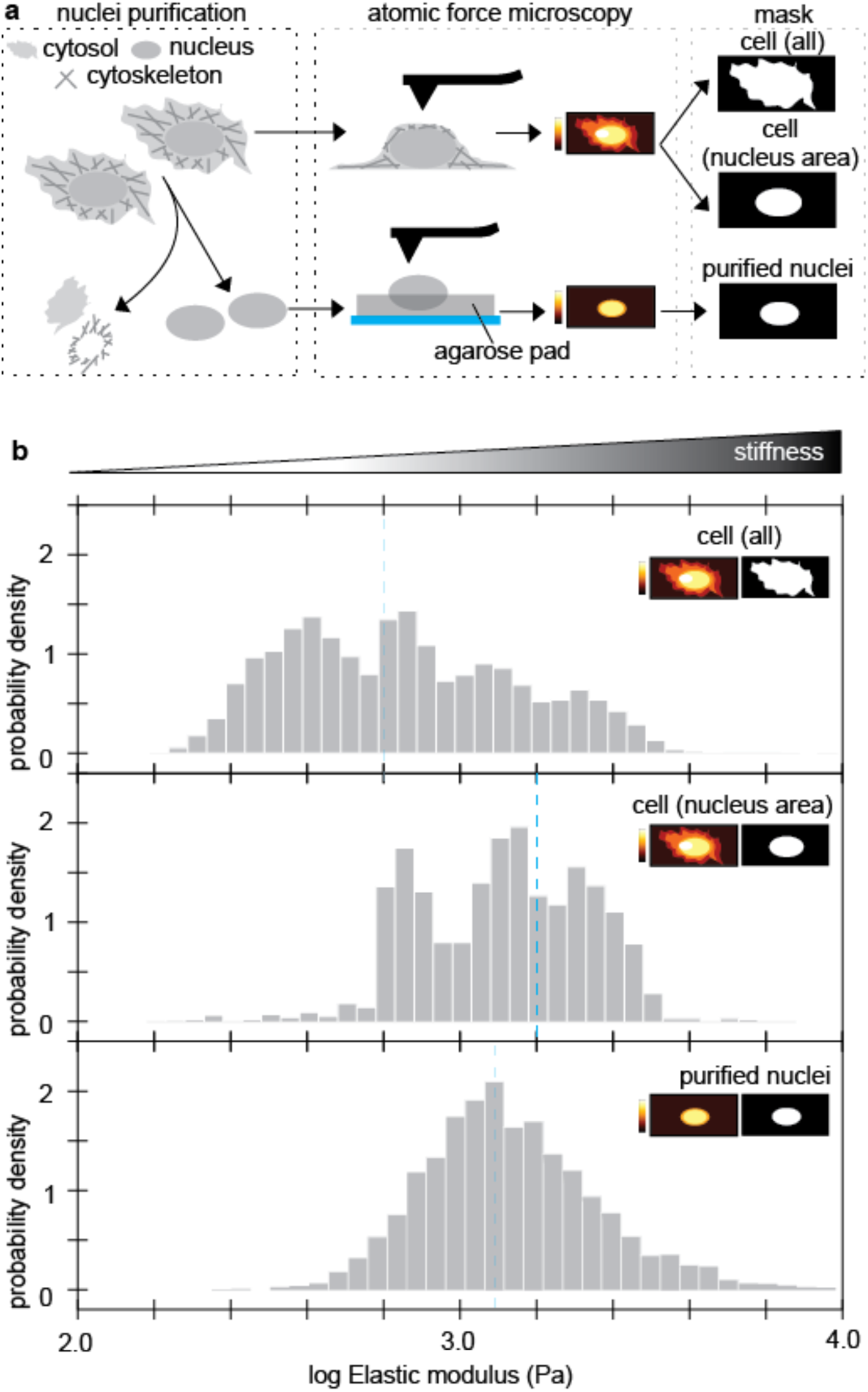
Nuclear stiffness does not depend on the cytosolic components. a. Scheme of the experiment. C2C12 myoblast nuclei were extracted to remove the cytosolic cytoskeleton. Atomic force microscopy force maps were acquired for the cells and the purified nuclei, the latter seeded in an agarose gel. To analyze the data, a mask was used to depict the cell, the nucleus area of the cell or the purified nucleus. b. Elastic modulus distribution calculated from the force maps. Dashed lines: median value for elastic modulus.

**Table 1.** Summary of elastic modulus analysis.

| Cell line | Sample | Mean elastic modulus (Pa) | Median elastic modulus (Pa) |
| --- | --- | --- | --- |
| C2C12 | mb cell | 939.6 | 693.1 |
| C2C12 | mb cell (nucleus area) | 1,535.3 | 1,636.5 |
| C2C12 | mb nuclei | 1,636.5 | 1,384.5 |
| C2C12 | mb MeCP2+ low nuclei | 16,285.6 | 13,319.5 |
| C2C12 | mb MeCP2+ high nuclei | 25,607.0 | 23,452.5 |
| C2C12 | differentiated nuclei | 3,395.9 | 1,993.2 |
| J1 wt | ESC nuclei | 13,110.5 | 2,414.6 |
| J1 wt | -LIF day 7 (D7) nuclei | 11,863.5 | 1,822.6 |
| J1 wt | -LIF day 14 (D14) nuclei | 158,706.6 | 12,666.5 |
| J1 wt | -LIF day 21 (D21) nuclei | 19,903.3 | 7,970.8 |
| J1 MeCP2 KO | ESC nuclei | 17,307.4 | 2,829.0 |
| J1 MeCP2 KO | -LIF day 7 (D7) nuclei | 2,242.8 | 821.8 |
| J1 MeCP2 KO | -LIF day 14 (D14) nuclei | 16,896.9 | 1,450.4 |
| J1 MeCP2 KO | -LIF day 21 (D21) nuclei | 19,903.3 | 7,970.8 |
| J1 NSC wt | NSC nuclei | 418,239.0 | 23,117.5 |
| J1 NSC wt | neuron nuclei | 317,223.6 | 20,697.0 |
| J1 NSC MeCP2 KO | NSC nuclei | 77,537.8 | 3,556.0 |
| J1 NSC MeCP2 KO | neuron nuclei | 94,031.6 | 18,055.0 |
| C2C12 | untransfected (mb) nuclei | 1,325.2 | 1,346.4 |
| C2C12 | + pMeCP2 wt nuclei | 2,929.1 | 2,258.8 |
| C2C12 | + pMeCP2 P101H nuclei | 1,081.1 | 970.8 |
| C2C12 | + pMeCP2 R106W nuclei | 1,243.9 | 1,032.1 |
| C2C12 | + pMeCP2 R133C nuclei | 1,307.9 | 1,277.7 |
| C2C12 | + pMeCP2 A140V nuclei | 2,936.4 | 2,526.2 |
| C2C12 | +pMeCP2 T158M nuclei | 2,936.4 | 2,526.2 |
| C2C12 | +pMeCP2 R168X nuclei | 1,320.0 | 1,073.5 |
| C2C12 | +pMeCP2 R255X nuclei | 1,911.7 | 1,967.5 |
| C2C12 | +pMeCP2 R270X nuclei | 1,619.5 | 1,755.0 |
| C2C12 | +pMeCP2 R270X nuclei | 2571.6 | 2,211.7 |

### MeCP2 dependent heterochromatin clustering increases stiffness

After validating the extraction of nuclei for AFM measurements, we aimed to further investigate the impact of heterochromatin on nuclear stiffness. To achieve this, we took advantage of a method we developed that allowed us to identify cell fractions with distinct, quantified levels of MeCP2 (*42*). This was accomplished through fluorescent-activated cell sorting (FACS) of myoblast transfected with a plasmid encoding GFP-tagged MeCP2. The rationale for using myoblasts is because, being adult stem cells, the endogenous level of MeCP2 in these cells is low to undetectable. MeCP2 reorganized the heterochromatin in a concentration dependent manner by clustering the heterochromatin compartments (Figure 2a), which led to a reduction in their number (Figure 2b) and an increase in their size (Figure 2c), consistent with previous findings in these cells (*33*, *35*). Nanomechanical characterization of the nuclei using AFM revealed an increase in the nuclear stiffness that correlated with the higher MeCP2 concentration (Figure 2d, Table 1). The median elastic modulus increased from 1.4 kPa in untransfected myoblast (containing ∼0.4 µM MeCP2 (*42*)) to 13.3 kPa in the low MeCP2 level fraction (∼11.8 µM MeCP2 (*42*)) and 23.5 kPa in the high MeCP2 level fraction (∼131.2 µM MeCP2 (*42*)).

**Figure 2.**
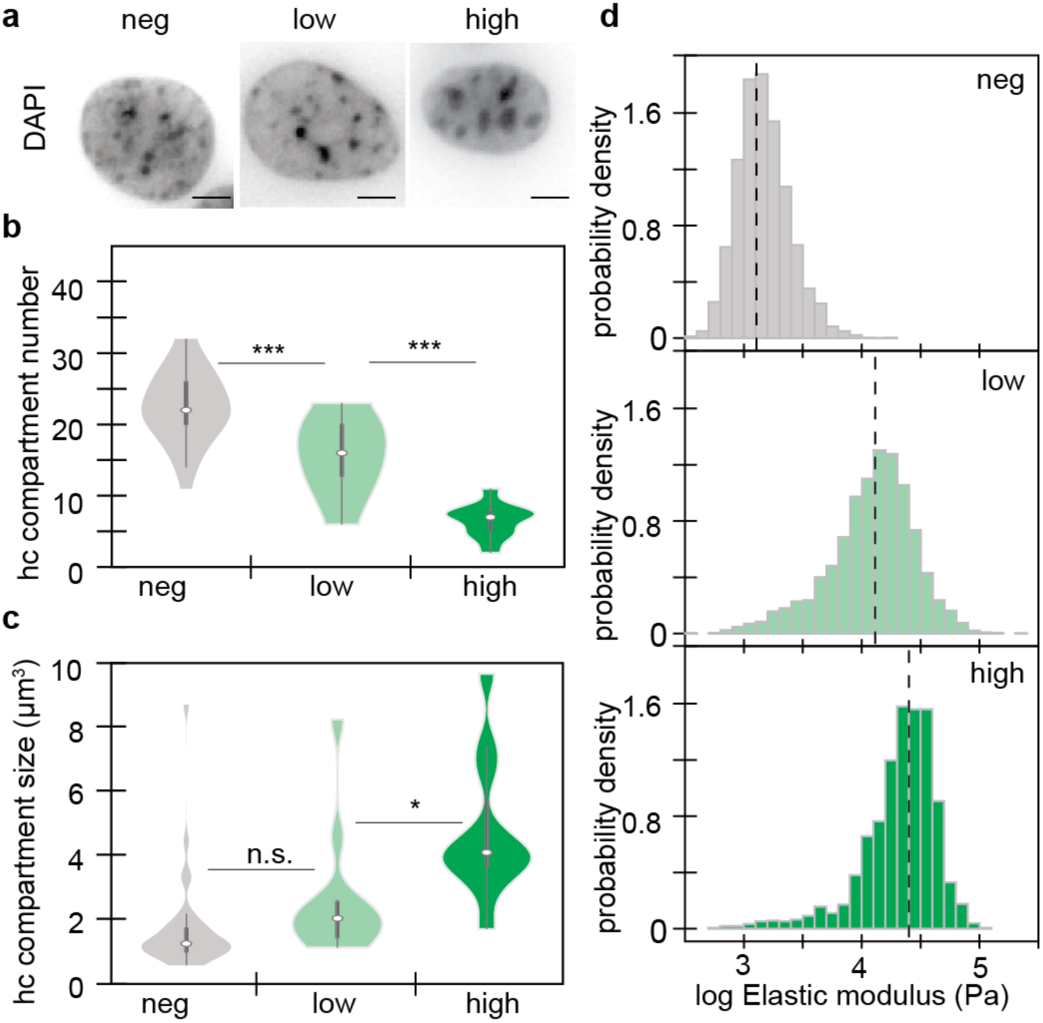
Heterochromatin clustering driven by MeCP2 increases nuclear stiffness. a. Visualization of the heterochromatin compartments by DAPI staining (high density foci). Scale: 5 µm. b-c. 3D quantification of the number (b) and volume (c) of the heterochromatin compartments for the different levels of MeCP2 in 10-20 nuclei. n.s.: non significant; *: p-value < 0.05; ***: p-value <0.0001. d. Elastic modulus profile of at least 30 nuclei from the different fractions based on MeCP2 levels separated by fluorescence activated cell sorting. Dashed line: median value.

### MeCP2 plays a major role in increasing the nuclear stiffness during neural differentiation

In view of the role of MeCP2 in neurological disease, we investigated whether nuclear stiffness is affected during neuronal differentiation. Therefore, we used a well-characterized model of embryonic stem cells (ESC) differentiation into neurons via LIF deprivation (Figure 3a). In this model, it is known that: i) MeCP2 level emerges after 14 days of differentiation (*34*); ii) significant differences in heterochromatin clustering exist between MeCP2 wild-type (wt) and MeCP2 knockout (KO) cells (*34*); and iii) cellular stiffness increases during differentiation (*43*). To study this, we generated and characterized MeCP2 KO ESC using a MIN-tag strategy (Figure S1a-d). We confirmed the loss of pluripotency in both wt and KO ESCs through immunostaining and high-throughput microscopy using the pluripotency marker Oct3/4 (Figure S1e-f), as well as the kinetics and absence of MeCP2 in wt and MeCP2 KO cells, respectively (Figure S1i,l). Moreover, we observed an increase in the neuronal marker NeuN during differentiation (Figure S1k). Notably, wt and MeCP2 KO cells were indistinguishable based on Oct3/4 and NeuN at each differentiation stage (Figure S1f,k). However, consistent with previous findings (*44*), MeCP2 KO nuclei were slightly larger than their wt counterparts (Figure S1i). We also confirmed impaired heterochromatin clustering in MeCP2 KO cells (*34*), with an increased number (Figure 3b) and reduced size (Figure 3c) of heterochromatin compartments, particularly at differentiation days 14 and 21, when MeCP2 level is highest in wt cells. Then, we purified the nuclei at every stage and derived the elastic modulus distribution from AFM measurements (Figure 3d, Table 1). In ESC, the median elastic modulus was similar between wt (2.4 kPa) and MeCP2 KO (2.8 kPa). However, upon differentiation, wt nuclei became significantly stiffer than MeCP2 KO counterparts with median elastic modulus as follows: 1.8 versus 0.8 kPa at day 7, 12.7 versus 1.5 kPa at day 14, and 28.4 versus 8.0 kPa in neurons.

**Figure 3.**
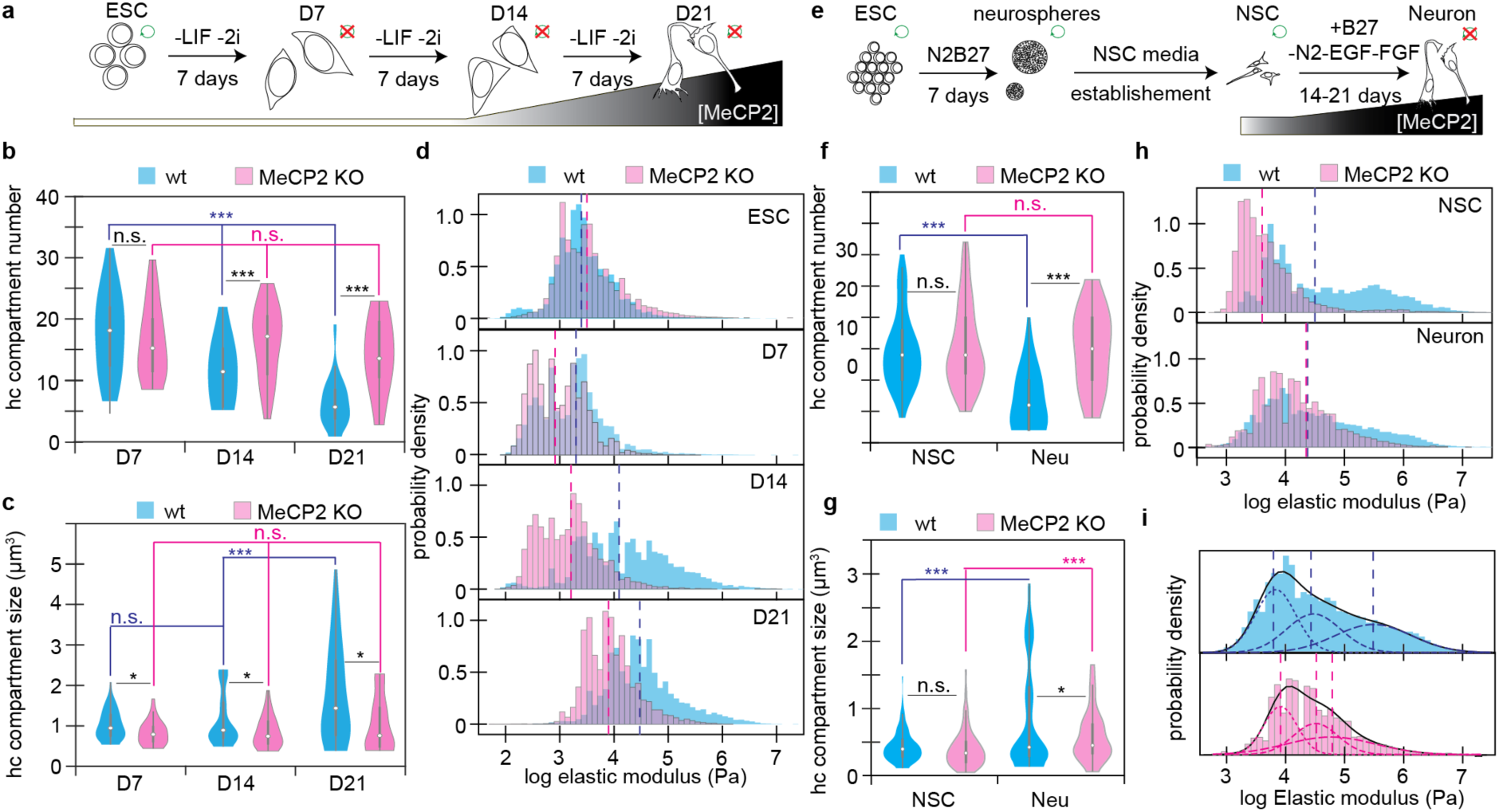
Deletion of MeCP2 severely impairs the increase of stiffness during neuronal differentiation. a. Differentiation model from embryonic stem cells (ESC) to neurons by leukemia inhibitory factor (LIF) deprivation, with overview to the renewal abilities (green circle) and the MeCP2 levels after 7, 14 and 21 days (D7, D14 and D21 respectively). b-c. 3D analysis of DAPI images and quantification of the number of heterochromatin compartments (b) and their volume (c) in the neural differentiation from ESC to neurons for wt (blue) and MeCP2 KO (rosa). d. Elastic modulus profile based on AFM measurements for the different stages of the differentiation of ESC to neurons (D21) in wt (blue) and MeCP2 KO (rosa) cells. Dashed lines: median. e. Differentiation model from ESC to neurons by generation of stable neural stem cells (NSC), which maintain the self-renewal ability (green circle) and the levels of MeCP2. f-g. 3D analysis of DAPI images and quantification of the number of heterochromatin compartments (b) and their volume (c) in the neural differentiation from NSC to neurons for wt (blue) and MeCP2 KO (rosa). h. Elastic modulus profile based on AFM measurements in NSC and neurons of wt (blue) and MeCP2 KO (rosa) cells. Dashed lines: median. i. Gaussian Mixture model showing 3 populations in the AFM measurements for neurons shown in h. The black lines represent the overall model, with the individual populations shown inside. The mean of each population is represented by vertical dashed lines. Note that the Y axes are adjusted for better visibility. n.s.: non significant; *: p-value < 0.05; ***: p-value < 0.0001.

To exclude differences in the differentiation timing at the precursor stage, we generated stable neural stem cells (NSCs) from neurospheres (Figure 3e, Figure S2a), which would be equivalent to the 14-day differentiation stage. MeCP2 KO NSCs showed slightly reduced Pax6 marker level (Figure S2b-c), but identical renewal ability based in the 5-ethynyl-2′-deoxyuridine (EdU) uptake (Figure S2d) and capacity to produce neurons (Figure S2e), despite lacking MeCP2 expression (Figure S2h). In some NSCs MeCP2 could be detected (Figure S2h) but the overall levels were rather low compared to the levels reached in differentiated neurons (Figure S2i). Like the previous ESC differentiation, MeCP2 KO NSCs and neurons had larger nuclei compared to wt counterparts (Figure S2f). Yet, heterochromatin clustering differences between wt and KO cells were only prominent at the neuron stage, as both wt and MeCP2 KO NSC cells displayed similar numbers (Figure 3f) and volumes (Figure 3g) of heterochromatin compartments.

AFM measurements from the nuclei showed approximately an order of magnitude difference in the median elastic modulus between wt and MeCP2 KO NSCs (Figure 3h, Table 1), with values of 23.1 kPa versus 3.6 kPa, respectively, whereas in the neurons, the median elastic modulus was only slightly higher in the wt with values of 20.7 kPa versus 18.1 kPa. However, when analyzed closely, the distribution of the elastic modulus was not equivalent and a Gaussian Mixture Model analysis revealed that the MeCP2 KO neurons had less of the stiffest population compared to the wt counterparts (Figure 3i), thus indicating a failure in increasing nuclear stiffness. In conclusion, our findings demonstrate that MeCP2 level plays a crucial role in controlling the increase in stiffness during neural differentiation.

### MeCP2-dependent increase of nuclear stiffness is compromised in mutations linked to Rett syndrome

We investigated whether MeCP2’s role in increasing nuclear stiffness is altered in the neurological disorder Rett syndrome. Therefore, we selected mutations and truncations that are known to cause the disease, including the most common ones (R106W, R133C, R168X, R255X, R270X, and R294X, Figure 4a) (*45*–*47*). We also included two mutations (P101H and A140V, Figure 4a) due to their unique heterochromatin-binding and clustering phenotypes (*48*): P101H binds like the wt, but fails to cluster heterochromatin, while A140V binds more strongly than the wt and forms larger, irregular clusters of heterochromatin.

**Figure 4.**
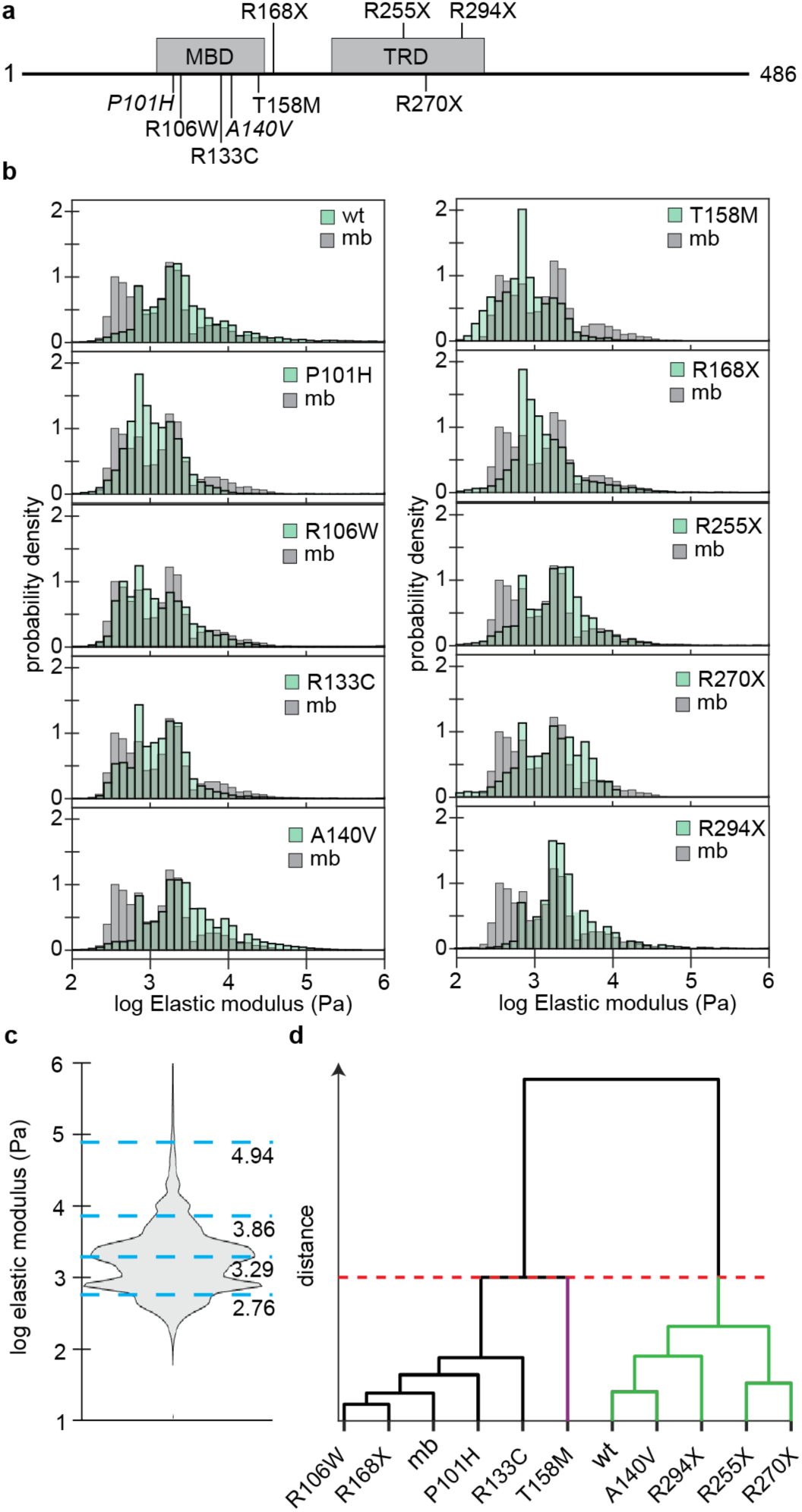
Rett syndrome mutations of MeCP2 impair the increase of nuclear stiffness. a. Scheme of the main structures of the MeCP2 protein (MBD: methyl binding domain; TRD: transcription repression domain) and the location of the mutations studied. b. Elastic modulus profile of the MeCP2 wt and mutations (green) versus the untransfected counterpart (mb, gray). c. Distribution of the elastic modulus for all conditions tested. Blue lines represent the centroids selected for k-means used for the clustering analysis, with the mean value stated underneath in log of Pa. d. Cluster analysis based on the percentages for each population in a 4-population k-means distribution.

Using the myoblast cells as a screening model, we transfected cells with plasmids encoding either wt GFP-tagged MeCP2 or mutant GFP-tagged MeCP2 and selected those that showed 50-70% transfection efficiency before purifying the nuclei for AFM measurements (Figure 4b, Table 1). The median elastic modulus was 1.3 kPa for the untransfected and 2.2 kPa for the wt MeCP2. Among the mutants, several exhibited median elastic modulus values within the range of untransfected cells and wt MeCP2, including R133C (1.3 kPa), R270X (1.8 kPa), R255X (2.0 kPa), and R294X (2.2 kPa). In contrast, some mutants demonstrated reduced stiffness compared to untransfected cells, with values of 1.1 kPa for R106W, 1.0 kPa for R168X, 0.9 kPa for P101H, and 0.8 kPa for T158M. Notably, the A140V mutation resulted in a higher elastic modulus of 2.9 kPa, surpassing that of wt MeCP2.

Given the differences in elastic modulus distributions among mutants, we used k-means clustering to group the data into four consistent populations with elastic moduli of 0.6, 1.9, 7.2, and 87.1 kPa (Figure 4c), based on the distribution of all the data collected. Cluster analysis (Figure 4d) grouped the mutants into three categories: one group included four mutations (P101H, R106W, R133C, and R168X) that aligned with the properties of untransfected cells. A second group comprised four other mutations (A140V, R255X, R270X, and R294X) that exhibited stiffness similar to wt MeCP2. Finally, the T158M mutant formed a distinct third group, characterized by its unique effect on nuclear stiffness.

These findings suggest that the impact of Rett syndrome mutations on nuclear stiffness can vary, ranging from a mild or full increase in stiffness (wt-like group), an impaired stiffness increase (untransfected-like group) to significant nuclear softening (mutant T158M).

### Stiffness changes related to MeCP2 are not linked to general changes in mechanobiology gene expression

Since chromatin reorganization is a known mechanism for altering gene expression, we explored whether the changes observed in the absence of MeCP2 or due to mutation could stem from alterations in the expression of genes that regulate the mechanical properties of the cell. For this purpose, we analyzed RNA-seq datasets from relevant samples, including brain cortex from MeCP2 KO models, excitatory neurons harboring R106W and T158M. An analysis of global gene expression changes associated with MeCP2 absence or mutation revealed a balance of upregulated and downregulated genes across all datasets (Figure 5a-c), indicating a dual role of MeCP2 as both a transcriptional activator and repressor.

**Figure 5.**
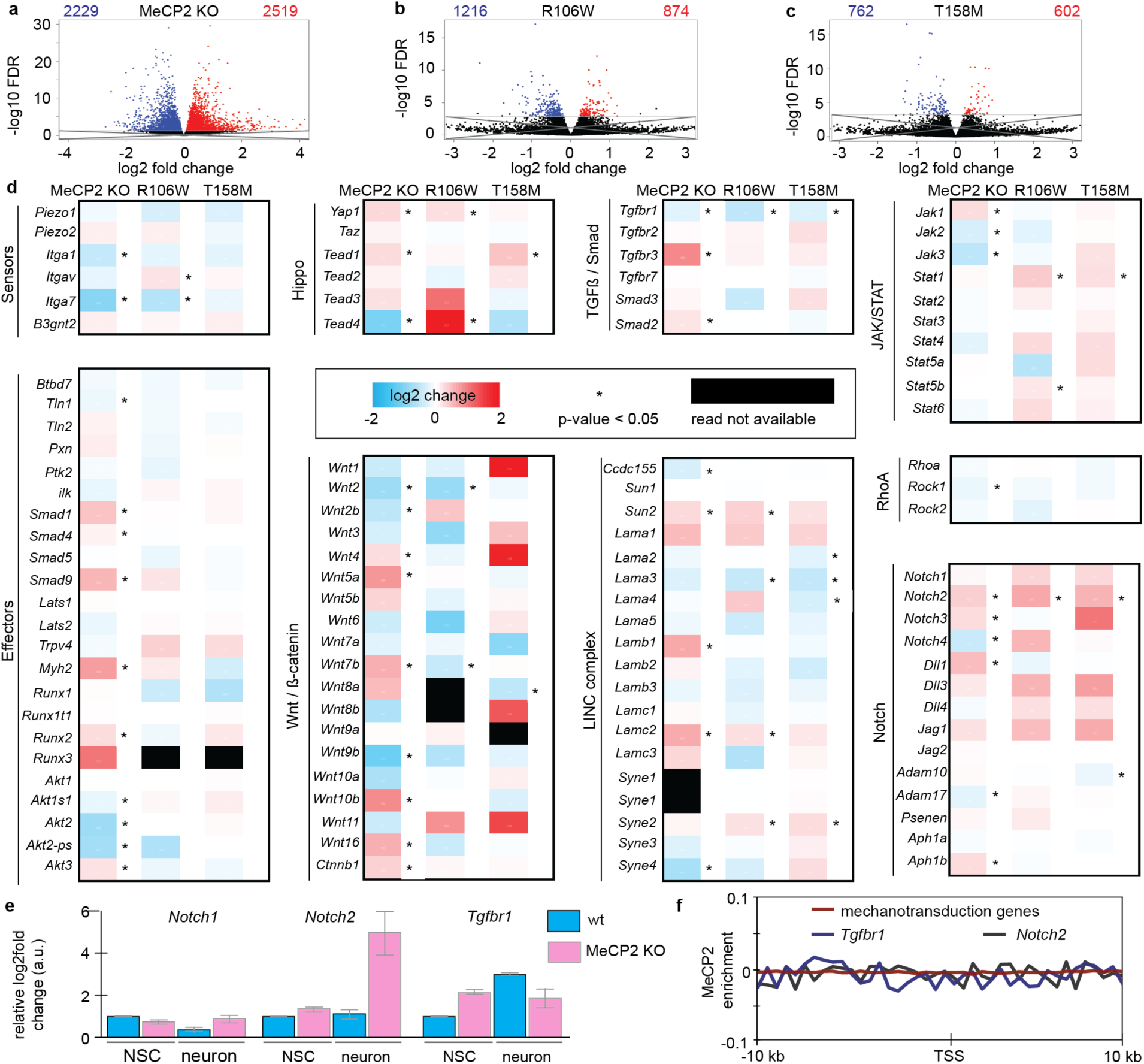
MeCP2 regulates the expression of the mechanotransduction-related genes *Notch2* and *Tgfbr1*. a-c. Volcano plots displaying the changes in overall expression of genes by the deletion of MeCP2 (a), or the mutation R106W (b) or T158M (c). Blue and red represent down- and up-regulated genes respectively with a false discovery rate (FDR) > 0.05. d. Gene arrays for the main components of the mechanotransduction pathways and its change due to MeCP2 deletion or mutation. e. Analysis of the expression of relevant genes in qRT-PCR in NSC differentiation. Bar plot shows the average and lines the standard deviation of three biological replicates, each of them done with three technical replicates. f. Chromatin immunoprecipitation analysis of MeCP2 in 20 kb around the transcription start site (TSS) of the mechanotransduction genes shown in d, as well as specifically for *Tgfbr1* and *Notch2*.

We next focused on genes implicated in mechanotransduction pathways, including sensors, receptors, transductors and effectors (Figure 5d). Interestingly, among these, only two genes showed significant and consistent changes across all conditions tested: *Tgfbr1* encoding for TFG-ß receptor I (TGFBR1), and *Notch2* encoding for Notch2 (Figure 5d). These changes were validated using reverse transcription followed by quantitative polymerase chain reaction (RT-qPCR) (Figure 5e).

We further assessed the dynamics of these genes in the differentiation models used. During the differentiation of wt NSCs, *Notch2* levels remained constant, however, in MeCP2 KO cells, its expression increased more than fourfold (Figure 5e). In contrast, Tgfbr1 mRNA levels increased during wt differentiation but remained unchanged in MeCP2 KO cells. Notably, at the NSC stage, Tgfbr1 levels in MeCP2 KO cells were already elevated compared to their wt counterpart (Figure 5e).

To confirm whether the observed changes in *Tgfbr1* and *Notch2* expression were directly driven by MeCP2 binding to their regulatory elements, we analyzed chromatin immunoprecipitation sequencing data of MeCP2 from mouse brain. The results showed no significant enrichment of MeCP2 binding, neither in the regulatory regions of mechanotransduction genes, nor in the regulatory regions of the two genes that were significantly changed (*Tgbfr1* and *Notch2*). These findings indicate that the changes in the expression of *Tgfbr1* and *Notch2* were not due to direct MeCP2 binding but rather arise from the reorganization of the heterochromatin caused by MeCP2 loss or mutation.

## Discussion

In this work, we demonstrated that MeCP2 changes the physical and mechanical properties of the nucleus through its heterochromatin clustering ability. Specifically, an increased MeCP2 concentration correlates with increased nuclear stiffness (Figure 2). This correlation extends to differentiation systems where MeCP2 levels rise, such as neural differentiation (Figure 3). Furthermore, we revealed that these changes in the physical properties are disrupted by mutations that lead to Rett syndrome (Figure 4). Finally, we showed that these alterations were not a consequence of widespread dysregulation of mechanotransduction genes (Figure 5), but rather the direct result of chromatin reorganization orchestrated by MeCP2.

The nucleus is the largest and stiffest organelle within the cell, playing a critical role in cellular mechanics. Studies have identified several nuclear components crucial to these mechanical functions, including nuclear membrane (*49*, *50*), lamin (*23*, *51*) and nuclear actin (*52*). While studies acknowledged chromatin as a contributor to nuclear stiffness (*52*), it is often considered in the context of its interaction with lamins (*51*, *53*). Here, we demonstrated that MeCP2-driven heterochromatin clustering (Figure 2b-c) alone is sufficient to enhance nuclear stiffness (Figure 2d), consistent with previous reports showing that increased heterochromatin density leads to a stiffer nucleus (*54*). We propose that the ability of MeCP2 to interact strongly with heterochromatin, as well as to self-associate, facilitates crosslinking that progressively increases the elastic modulus until either saturation of the binding sites or steric hindrance is reached, thereby limiting further compaction and stiffness.

Studies on the relevance of nuclear stiffness in cellular function have focused on two major points: mechanical stress adaptation and mechanoreception. Mechanical stress adaptation (in particular in muscle) is mostly dependent on the laminA concentration, as its levels increase with tissue stiffness (*11*). Mechanoreception also relies on the mechanical properties of the nucleus for cellular proprioception and mechanotransduction. In proprioception, the physical stimuli applied to cells are only sensed upon nuclear deformation (*2*). In mechanotransduction, the import of key transcription factors to the nucleus is controlled by nuclear stiffness. For instance, factors such as YAP are imported into the nucleus only when the mechanical stimuli are recognized by the nucleus (*55*, *56*). In the brain, where laminA levels are low (*11*), we found that chromatin organization determines nuclear stiffness (Figure 3d) rather than being only a response to mechanical stress (*19*, *57*, *58*). Neurons, which have higher MeCP2 levels than glial cells (*59*, *60*), may utilize this mechanism to elicit distinct responses than glial cells to identical mechanical stimuli.

The temporal dynamics of MeCP2 expression during brain development have been well-documented (*61*–*63*). In our investigation into the role of MeCP2 in nuclear stiffness, we identified two genes, Notch2 and Tgfbr1, that exhibited significant changes in MeCP2 KO mice (Figure 5e-f). Both factors are implicated in neuronal maturation. The Notch signaling pathway is downregulated during neurogenesis (*64*), with Notch2 specifically reported to inhibit neuronal differentiation (*65*, *66*). In contrast, TGFBR1 is active in neurons, and its downregulation reduces survival and maturation of the newborn neurons during neurogenesis

(*67*). Intriguingly, TGFBR1 is upregulated in other Rett mutations, such as R255X (*68*), which may contribute to the previously reported deficits in neurite maturation observed in MeCP2 KO and Rett syndrome (*69*, *70*). Multiple studies have demonstrated interactions between Notch2 and TGF-β signalling (*71*–*73*), Primarily in the regulation of cell fate determination and extracellular matrix remodeling (*74*–*77*). Although we cannot exclude a possible role of these proteins in modulating nuclear mechanical properties, there is currently no direct evidence supporting such an effect. Instead, our findings suggest that chromatin organization may play a more prominent role in determining nuclear stiffness.

Our analysis of Rett-associated MeCP2 mutations revealed a correlation between nuclear stiffness and disease severity (*78*, *79*). Mutations clustering with wt MeCP2 generally aligned with milder phenotypes, while those deviating from the wt cluster correlated with more severe cases (Figure 4c). The only exceptions for these correlations are R133C (distant from wt but mild) and R255X (close to wt but severe). This would explain why most of the Rett variants of MeCP2 are partially deficient in reorganizing the heterochromatin (*48*). Interestingly, our results showed that the mutant T158M has a softening effect on nuclear stiffness (Figure 4b-c). This specific mutation is associated with a severe clinical phenotype (*79*, *80*), but, until now, has shown limited impact in traditional *in vitro* assays (*48*, *81*, *82*). Further investigation on this specific mutation regarding the effect on the nuclear mechanics could provide critical insights into the mechanisms underlying Rett syndrome and inform therapeutic strategies.

## Methods

### Cell culture conditions

All cells used were deemed mycoplasma free. A list with the main characteristics, as well as reference to the original publications of the cell lines, can be found in Table S2.

Murine C2C12 myoblasts were grown in Dulbecco’s Modified Eagle’s medium (DMEM, Cat. No.: 41965039, Gibco) high glucose supplemented with 20% fetal bovine serum (Cat. No.: FBS-22A, Capricorn Scientifics), 110 mg/l sodium pyruvate (Cat. No.: 113-24-6, Sigma Aldrich), 1x L-glutamine (Cat. No.: G7513, Sigma Aldrich) and 1 µM gentamicin (Cat. No.: G1397, Sigma Aldrich), at 37 °C in a humidified atmosphere with 5% CO_2_. For passaging, growth media was aspirated, and cells were briefly washed with 0.02% ethylenedinitrilotetraacetic acid (EDTA, Cat. No.: A5097, AppliChem GmbH) in phosphate buffer saline (PBS), composed of 137 mM NaCl (Cat. No.: 3957, Carl Roth), 2.7 mM KCl (Cat. No.: P9541, Sigma Aldrich), 1 mM Na_2_HPO_4_ · 7 H_2_O (Cat. No.: X987, Carl Roth) and KH_2_PO_4_ (Cat. No.: 3904, Carl Roth), before incubation with trypsin-EDTA (Capricorn Scientific, TRY-3B). After 5 min at 37 °C, trypsin was inactivated by the addition of 2× volumes of growth media. In the case of transfection, cells were centrifuged at 1,400 rpm for 5 min and resuspended in 100 µl AMAXA M1 solution containing 2-10 µg of plasmid. AMAXA M1 solution was composed of 5 mM KCl (Cat. No.: 7447-40-7, Sigma Aldrich), 15 mM MgCl_2_ · 6 H_2_O (Cat. No.: 7786-30-3, Sigma Aldrich), 120 mM of Na_2_HPO_4_/NaH_2_PO_4_ (Cat. No.: 7558-79-4, Sigma Aldrich), and 50 mM mannitol (Cat. No.: 69-65-8, Caesar & Lorentz). Then, cells were electroporated using AMAXA nucleofection system (Lonza, S/N; 10700731), program B-032.

Murine J1 embryonic stem cells (ESC) were grown on feeder-free gelatine-coated culture dishes at 37 °C in a humidified atmosphere with 5% CO_2_. The coating of dishes or coverslips, when necessary, was performed by incubation in 0.2% gelatin from porcine skin (Cat. No.: G2500, Sigma Aldrich) in H_2_O for 15 min at room temperature. ESC growth media consisted of DMEM high glucose supplemented with 15% fetal bovine serum, 1× MEM non-essential amino acid solution (Cat. No.: M7145, Sigma Aldrich), 1× penicillin/streptomycin (Cat. No.: P0781, Sigma Aldrich), 1× L-glutamine (Cat. No.: G7513, Sigma Aldrich), 0.1 mM beta-mercaptoethanol (Cat. No.: 4227, Carl Roth), 1000 U/ml recombinant mouse leukemia inhibitory factor (LIF), 1 µM PD032591 (Cat. No.: 1408, Axon Medchem) and 1 µM CHIR99021 (Cat. No.:1386, Axon Medchem). Media was changed every day. For passaging, growth media was removed, and cells were briefly washed with PBS before incubation with trypsin-EDTA solution for 5 min at 37 °C. After the incubation, trypsin was inhibited by the addition of 2× volumes of ESC growth media. In case of transfection, cells were centrifuged at 1,400 rpm for 5 min and resuspended in 100 µl AMAXA M1 solution containing 2-10 µg of plasmid and electroporated using AMAXA nucleofection system, program A-023.

J1 ESC differentiation was performed by LIF and inhibitors deprivation supplemented with retinoic acid as described previously(*34*, *83*). Briefly, 10^3^ cells/cm^2^ were seeded and incubated overnight in ESC growth media. Then, media was removed, cells were briefly washed with PBS, and ESC differentiation media was added. ESC differentiation media had the same composition of ESC growth media, apart from LIF, PD032591, and CHIR99021, which were removed, and the addition of 10 µM retinoic acid (Cat. No.: R2625, Sigma Aldrich). Cells were then incubated at 37 °C and 5% CO_2_ for 7, 14, or 21 days, changing the media every second or third day.

Murine J1 neural stem cells (NSC) were derived from J1 ESC and established as described in the next section. NSCs were grown in plates, slide chambers, or coverslips coated with poly-D-lysine and laminin, prepared in advance, and stored at -20 °C. For coating, plates, slide chambers or coverslips were incubated in 10 µg/ml poly-D-lysine (Cat. No.: P7405, Sigma Aldrich) in H_2_O for 4 h and dried at room temperature for 20 min before an overnight incubation at 37 °C with 5 µg/ml laminin (Cat. No.: L2020, Sigma Aldrich) in ice-cold DMEM-F12 medium (Cat. No.: 56498C, Sigma Aldrich). NSCs were grown at 37 °C in a humidified atmosphere with 5% CO_2_ in NSC growth media, composed of Euromed-N (Cat. No.: ECL-ECM0883L, Biozol Diagnostica Vertrieb), 1× N-2 supplement (Cat. No.: 17502048, ThermoFisher Scientific), 1× L-glutamine (Cat. No.: G7513, Sigma Aldrich), 1× penicillin/streptomycin (Cat. No.: P0781, Sigma Aldrich), 20 ng/ml murine epidermal growth factor (Cat. No.: 315-09-500UG, Peprotech) and 20 ng/ml murine fibroblast growth factor-2 (Cat. No.: PPT-450-33-500, Peprotech). For passaging, media was removed, and cells were briefly washed with PBS before incubation with accutase (Cat. No.: A6964, Sigma Aldrich) for 3 min. Accutase was then inactivated by the addition of 2× volumes of NSC growth media. For J1 NSC differentiation to neurons, 2.5×10^4^ cells/cm^2^ were seeded in slide chambers or 60 mm plates coated as described above and cultured in NSC growth media for 48 h. Then, the media was changed to neuronal differentiation media composed of 3:1 Gibco Neurobasal media (Cat. No.: 21103049, ThermoFisher Scientific) and DMEM-F12 (Cat. No.: 56498C, Sigma Aldrich), 0.5x N-2 supplement (Cat. No.: 17502048, ThermoFisher Scientific)), 1x Gibco B-27 supplement (Cat. No.: 12587010, ThermoFisher Scientific), 1× penicillin/streptomycin (Cat. No.: P0781, Sigma Aldrich), 1x L-glutamine (Cat. No.: G7513, Sigma Aldrich), 10 ng/ml murine fibroblast growth factor-2 (Cat. No.: PPT-450-33-500, Peprotech), 20 ng/ml brain derived neurotrophic factor (BDNF, Cat. No.: AF-450-02-10UG, Peprotech). Differentiation media was changed every 3-4 days for 14-21 days to obtain differentiated neurons.

### Generation of J1 ESC MeCP2 KO cell line

To generate J1 ESC knockout for the *Mecp2* gene (MeCP2 KO), a MIN strategy was followed (*84*). First, a J1 ESC containing the MIN tag in the exon 2 of the *Mecp2* gene (J1 ESC MIN-MeCP2) was generated by CRISPR/Cas9 (Figure S1a). To do so, a plasmid containing the template for the guide RNA (CACCGTCAGAAGACCAGGATCTCCA) as well as the Cas9 was generated. DNA oligos (forward and reverse) coding for the guide RNA (Table S2) were annealed by mixing 100 µM in NEB 4 buffer, incubated at 95 °C for 5 min, and then cooled at room temperature for 10 min. The annealed DNA, together with the pSpCas9(BB)-2A-Puro (Addgene #62988, Table S3), was incubated with BpiI (ThermoFisher Scientific) and 30 U T4 DNA ligase (Cat. No.: M0202, New England Biolabs) in T4 DNA ligase buffer, in 55 cycles of 5 min 37 °C and 5 min 20 °C followed by 60 min at 37 °C and 10 min at 95 °C and the mix used to transform TOP10 *Escherichia coli* (Cat. No.: C404003, ThermoFisher Scientific). The resulting plasmid was verified by DNA sequencing. J1 ESC were then transfected with the previously constructed plasmid together with the single-stranded repair template containing the MIN tag, which was chemical synthetized (CTTCTTTGTCCTCCTTCTTGTCTTTCTTCGCCTTCTTAAACTTCAGTGGCTTGTCTCTGAGGCCCTGGAGATCCTGGGTTTGTACCGTACACCACTGAGACCGCGGTGGTTGACCAGACAAACCGTCTTCTGACTTTTCCTCCCTGAAGTATTAAACAAATATGTAAGTATTACAGAGAACACAGCTGTCTGCACAGTAG). Cells were seeded onto 365-well plates for selection with 10 µg/ml puromycin (Cat No.: ant-pr-1, InvivoGen) for 2 days. The presence of the MIN tag was then checked by genomic DNA isolation followed by PCR (Figure S1c) with the oligos described in Table S2 and the DNA from positive clones was sequenced to confirm the correct MIN tag integration in exon 2 of the *Mecp2* gene locus (Figure S1c).

Subsequently, the MeCP2 KO was generated by introducing a stop codon in the MIN tag. This was achieved by transfecting the J1 ESC MIN-MeCP2 with a plasmid coding for the Bxb1 recombinase (pCAG-NLS-Bxb1, Addgene #65625, Table S3) together with a plasmid containing the attB site, followed by mCherry cDNA and a stop codon (pattB-Cherry-Stop-puro, addgene #65529, Table S3). Transfected cells were grown in 365-well plates and selected by genomic DNA screening PCR (Figure S1d). The cell line generated was further characterized by verifying its ability to form colonies (Figure S1b), its pluripotency by staining for the Oct4 marker (Figure S1e-f), its self-renewal by labeling and staining for proliferating cells with EdU (Figure S1g-h), as well as the ability to differentiate into neurons by NeuN staining (Figure S1k-l). We also confirmed by immunostaining the absence of MeCP2 protein during differentiation (Figure S1j and S1l).

### J1 NSC cell line generation

J1 ESC wt or J1 ESC MeCP2 KO were cultured at 37 °C in a humidified atmosphere with 5% CO_2_ in 0.2% gelatin-coated T25 flask in Gibco KnockOut Dulbecco’s modified Eagle’s medium (Cat. No.: 10829018, Thermo Fisher Scientific) supplemented with 15% KnockOut serum replacement (Cat. No.: 10828028, Thermo Fisher Scientific), 1× MEM non-essential amino acid solution, 1× penicillin/streptomycin (Cat. No.: P0781, Sigma Aldrich), 1× L-glutamine (Cat. No.: G7513, Sigma Aldrich), 0.1 mM beta-mercaptoethanol and 1000 U/ml recombinant mouse LIF. Cells were passaged multiple times by briefly washing with PBS and then incubated with accutase for 3 min at 37 °C until they grew as monolayers (Figure S2a, left panel). Then, 4×10^4^ cells/cm^2^ were seeded in 0.2% gelatin-coated 6-well plates and cultured in neurobasal media and DMEM-F12 (1:1) medium supplemented with 0.1 mM beta-mercaptoethanol, 0.5× N-2 supplement (Cat. No.: 17502048, ThermoFisher Scientific), 0.5× B-27 supplement (Cat. No.: 12587010, ThermoFisher Scientific), 1x GlutaMAX supplement (Cat. No.: 35050038, Thermo Fisher Scientific) and 1× penicillin/streptomycin (Cat. No.: P0781, Sigma Aldrich) for up to 7 days at 37 °C in a humidified atmosphere with 5% CO_2_ until maximal confluence was reached. Then, cells were dissociated with accutase, reseeded in non-coated T25 flask, and incubated in NSC growth media at 37 °C and 5% CO_2_ for 2-4 days allowing the formation of neurospheres of 100-200 µm diameter (Figure S2a, middle-left panel). Once they reached this diameter, neurospheres were transferred to a 15 ml tube using a wide bore tip and spinned down at 500 rpm, then resuspended gently in fresh NSC media. Then, neurospheres were seeded in poly-D-lysine/laminin-coated T25 flasks, where neurospheres were attached to the dish and started the differentiation to NSC. Once confluence in the T25 flask was reached, cells were passaged using accutase and split at 1:2 ratio into new culture dishes. Stable NSC cell lines were established after 15 passages (Figure S2a, middle-right panel). NSCs were validated using Pax6 marker immunostaining (Figure S2b-c) and their self-renewal ability was confirmed by labeling and staining for proliferating cells with EdU (Figure S2d). The absence of MeCP2 protein was again confirmed by immunostaining with anti-MeCP2 specific antibodies (Figure S2g).

### Cell immunostaining and EdU click-it reaction

For immunostaining, cells were grown on appropriately coated coverslips or slide chambers and differentiated (if required) as described above. Cells were washed twice with PBS and then fixed using either 3.7% formaldehyde for 10 min or ice-cold methanol for 6 min depending on the antibodies used as described in Table S4. After three times washing with PBS, cells were permeabilized with 0.5% Triton X-100 (Cat. No.: 10670, LS Laborservice) in PBS and washed three times with 0.01% Tween-20 (Cat. No.: 9127.1, Carl Roth) in PBS (TPBS). Then, cells were incubated in a blocking solution composed of 1% BSA in PBS for 20 min. After blocking, samples were incubated with primary antibody, undiluted or diluted in blocking solution as described in Table S4 for 2 h. In each immunostaining assay, a secondary control was performed following the same procedure but skipping the primary antibody incubation. Non-bound antibodies were washed three times with TPBS before adding the secondary antibody diluted in PBS as described in Table S4, followed by another three washing steps with TPBS. Samples were then counterstained with 1 µg/ml of the DNA dye DAPI, washed twice in PBS and once in distilled H_2_O, dried, and mounted on a slide into a drop of Mowiol mounting media composed of 13% Mowiol 4-88 (Cat. No.: 81381, Sigma Aldrich), 33% glycerol (Cat. No.: G9422, Sigma Aldrich), 2% 1,4-diazabicyclo-[2.2.2]octane (Cat. No.: D2522, Sigma Aldrich) and 133 mM Tris-HCl pH 8.5 (Cat. No.: 93362, Sigma Aldrich).

For EdU (Cat No: 7845.1, Carl Roth) incorporation detection, cells were grown on appropriately coated coverslips and pulsed with 10 µM EdU for 20 min. EdU can only be integrated into the genome during active DNA replication. A click-it reaction, following the manufacturer’s protocol, was performed to link the EdU with a dye: 6-carboxyfluorescein (Cat. No.: 7806.2, Carl Roth) or Eterneon-Red 645 (Cat. No.: 1Y73.1, Carl Roth). DNA was counterstained with 1 µg/ml of DAPI for 15 min and cells were mounted in a glass slide with Mowiol mounting media.

### Flow cytometry and GFP-intensity based categories

Cells were transfected as described above and one untransfected plate was grown in the same conditions. Cells were harvested 20 h after transfection, resuspended in PBS, and separated according to their transfection level by fluorescence-activated cell sorting (FACS) on a S3 Cell Sorter (Table S5) into three gates as described in (*35*, *42*). Briefly, cells were exposed to a 488 nm laser, and intensity was measured after a 525 ± 30 nm emission filter. Cells were plotted against log10 of the sum intensity. Intensities were grouped into bins with a value calculated as the difference between the maximum of the transfected and untransfected samples divided by 42. Then, gates were defined as follows: negative (values lower than 8×bin), low expressing (values within 13×bin and 22×bin), and high expressing (values within 24×bin and 33× bin).

### Nuclei purification

We modified a protocol described previously (*85*). Cells were dissociated from the plate by trypsin-EDTA solution and resuspended in the correspondent growth media. To remove the media, cells were centrifuged at 1,400 rpm for 5 min and the supernatant was discarded. Then, pellets were resuspended in ice-cold PBSN, composed of 0.1 % Nonidet P-40 substitutive (Cat. No.: 74385 Sigma Aldrich) in PBS, triturated by continuous pipetting, and pelleted by short centrifugation at high speed (13,000-16,000 rpm) for 20 seconds. When needed, pellets were then resuspended in PBSN containing 1 µg/ml DAPI and incubated at room temperature for 15 min or, alternatively, resuspended in PBSN without incubation time. A second centrifugation at high speed for 20 seconds was done and the resulting pellet was resuspended in buffer A2, composed of 20 mM Triethanolamine-HCl (Cat. No.: T-1377, Sigma Aldrich) buffer, 30 mM KCl, 10 mM MgCl_2_ • 6 H_2_O, 0.25 M sucrose (Cat. No.: 4661, Carl Roth) and 0.1 mM phenylmethylsulfonyl fluoride (Cat. No.: 6367, Carl Roth). In this buffer, nuclei were kept at 4 °C for a maximum of one week.

### Plasmids

All plasmids used and their characteristics are listed in Table S3.

The plasmids pEG-MeCP2 R270X and pEG-MeCP2 R294X were generated using a Q5 directed mutagenesis from the pc1208 (pEG-MeCP2) following the standard protocol described by the manufacturer (NEB, E554S). The primer pairs flanking the regions to be deleted are listed in Table S2. The mutation was added by DNA polymerization reaction, and the reaction product was then incubated in a buffer with kinase, ligase and DpnI to phosphorylate and ligate the newly generated plasmid containing the mutation while digesting away the template. Finally, *E. coli* TOP10 cells were transformed with the reaction mix. Plasmids were then sequenced to confirm the mutation.

### Atomic force microscopy

For each group of experiments, nuclei were seeded in partially dehydrated agarose pads, composed of 0.5% agarose (Cat. No.: A9539, Sigma Aldrich) in PBS evenly distributed in 15 mm glass coverslips. Once the buffer in which the nuclei were resuspended was drained in the agarose, fresh buffer A2 was added on top during the measurements. The samples were selected using bright-field microscopy on the AFM setup to position the tip at regions containing islands of nuclei (with ≥5 nuclei in each frame).

We employed Force-Volume (F–V) mode, which involves static loading rather than oscillatory measurements. Consequently, the measured modulus reflects contributions from both shear and compressive deformation, though compressive effects are limited due to material incompressibility. Regardless of the time scale, local indentation captures both deformation types as the tip indents the material until the setpoint force is reached. The reported modulus is not an absolute measure but is used for comparative analysis of relative differences in nuclear mechanics under varying chromatin reorganization conditions. F–V mapping was carried out using a Nanowizard II AFM (JPK Instruments) coupled with a Zeiss Axio Observer Z1 optical microscope (Carl Zeiss). This technique employs a linear ramp method, where a complete force‒distance (F‒D) curve is recorded at each pixel. Soft cantilevers with nominal spring constants of 0.06 N/m (triangular-shaped microlevers, SNL-D, nonconductive sharp silicon nitride from Bruker) were employed for measuring the mechanical properties and performing spectroscopy of the cells. These cantilevers have a nominal flexural resonance frequency of 18 kHz in air and a sharp tip radius of 2 nm. The inverse optical lever sensitivity of the AFM system was determined by recording a single F‒D curve on the glass substrate (coverslip) of the nuclei in fresh buffer A2. Additionally, the cantilever spring constant was calibrated using the thermal noise method (*86*). For comparative analysis of the stiffness values obtained from the control measurements through F‒V mapping in this study, the following experimental parameters were set: the trigger point was adjusted to 2 nN to ensure a large indentation range (approximately 1000 nm), the z-length was set to 3 µm, the extension time was 90 s, the maximum sample rate was 16,000 Hz, and the grid size was 30 x 30 µm² with 64 x 64 points. The entire force-volume map was subsequently processed using MATLAB code to generate a stiffness distribution as demonstrated in our previous studies (*87*).

The F-D curve processing steps began with correcting the baseline tilt by fitting a line to the data between 50% and 99% of the z-piezo values (cantilever completely withdrawn at 100%). The contact point was identified as the z-piezo position where the cantilever was in further approach, and the force remained within the positive (repulsive) interaction regime. To calculate the tip indentation depth, the distance between the AFM tip and the sample surface was determined by subtracting the cantilever deflection from the z-piezo position. These force-indentation (F–d) curves were then fitted to Hertz contact mechanics models to produce local elasticity (E) maps, based on the following formula (Equation 1)

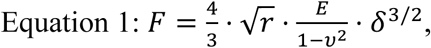

being F the force, r the radius of the curvature of the tip, 𝝂 the sample’s Poisson ratio (lateral to axial strain, ranges from -1 to 0.5), and δ is the indentation depth. The global elastic modulus values for each nucleus were obtained by averaging the cumulative local elastic modulus values, which were derived from 4096 F–D curves measured across the surface of nuclei.

### High throughput microscopy

Cells were immunostained as described above. For image acquisition, a Nikon-CREST widefield microscope was used (Table S5). During imaging, the acquisition conditions (exposure time, optical gain, and laser power) were kept constant between samples in the channel containing the protein of interest (POI), while the counterstaining channel (generally DAPI) conditions were modified to get the best segmentation of individual nuclei in each field.

Analysis of the images was done in ImageJ. First, the DAPI channel was used for segmenting the nuclei. Standard processing of Gaussian blur (sigma of 5 or 10 pixels if 20X or 40X objective was used respectively) followed by a background subtraction (rolling background of 20 or 50 pixels, in 20X or 40X images respectively) and Otsu auto-threshold. The nuclei were defined as ROI using the Analyze Particle function only restricting the edges and checked manually to remove or modify nuclei that were not properly segmented.

Once the ROIs in the nuclei were determined, the area and the total intensity (IntDen) was calculated for each channel. For comparing between conditions, intensity values from at least three biological replicates were merged and compared in a violin plot. Statistical differences were calculated using F-test assuming equal variance between samples.

### Acquisition and analysis of 3D samples

Z-stacks were taken following Nyquist sampling in a Leica TSC SPE-II confocal point scanner microscope (Table S5). For the 3D analysis using ImageJ 3D suite, individual cells were segmented manually and then processed to: i) get the nuclear volume, by applying a 3D median filter with 3x3x2 pixels (xyz) followed by Otsu threshold segmentation; ii) get the heterochromatin compartments, by using deconvolution as described (*88*). The resulting image was thresholded and applied a skeletonize to generate a seed that was then used for the 3D spot segmentation of the 3D suite, together with the deconvolved image. The resulting spots were then filtered by the nuclear mask. Volumes and numbers were then taken from the respective masks using 3D measurements of the 3D suite.

### RNA-seq and ChIP-seq

We used published datasets for the RNA-seq and ChIP-seq. For RNA-seq, the raw counts of genes in RNAseq analysis were downloaded from the GEO repository. We used the dataset GSE140054 (*89*) for the comparison of wt and MeCP2 KO, specifically the samples GSM4152139-GSM4152148, which correspond to cortex of wt and MeCP2 KO mice, 5 replicates each. Additionally, we used the dataset GSE83474 (*90*) to compare the wt MeCP2 versus the Rett mutations, using the samples GSM2203994-GSM2204004, which correspond to 4 replicates of excitatory neurons of a pool of 2-3 mice (six weeks-old) for Mecp2^wt/Y^, Mecp2^R106W/Y^ and Mecp2^T158M/Y^. The differential DESeq2 Plots were visualized using RStudio (R version 4.4.0, Version 2024.04.1+748 (2024.04.1+748)).

Sratoolkit (version 2.11.0) was used to download the ChIP-seq datasets from the GEO-database (Gene Expression Omnibus, https://www.ncbi.nlm.nih.gov/geo/), specifically the dataset GSE90704 (*91*) and the samples G2410973-GSM2410978, which correspond to brain cortex of six weeks-old wt mice. Reads quality was checked using the FastQC program (version 0.11.9). When necessary, Trimmomatic (version 0.36) was used to remove reads of poor quality. Next, bowtie2 (version 1.3.1), was used to align the reads to the mouse genome (mm10 genome assembly, https://hgdownload.cse.ucsc.edu/goldenpath/mm10/bigZips/) and Samtools (version 1.10) was used to generate bam files. Samtools were also used to merge the 3 replicates of MeCP2 Chip-seq/input correspondingly. The input signal was subtracted from the Chip-seq signal using deepTools (version 3.5.1_singularity), command bamCompare, using the option subtract. DeepTools were also used to plot profiles around the TSS of genes identified as unchanged/ changed in R106W and changed in T158M.

### Violin plots, statistics, and Gaussian mixture model

The representation of the data as violin plots, as well as the probability density functions (pdf), was done in MATLAB R2021a update 8 (6.10.0.2198249), using GAVI, a self-written script. The script is available in the TU-datalib (see data availability section). GAVI collects the data from a “.csv” (comma-separated values) file which contains the conditions in individual columns and represents them as individual probability density function graphs and/or violin plots. Additionally, it performs a Gaussian Mixture Model (GMM) analysis, in which the model with lesser Bayesian Information Criteria is selected as the best fitting, to divide the data into populations, providing the mean value of each population as well as the percentage of the total measurements that would fall into the respective populations. Lastly, it performs a two-sided t-test with the null hypothesis of equal means with equal variance between all the datasets to quantify the significance of differences between the data.

For the elastic modulus pdf representations, the size of the bars is 0.1 log (Pa). The use of logarithms in these graphs was chosen to enhance visibility, but neither the differences nor the GMM analysis changed when using the raw data.

In the violin plots, the x spread represents the frequency of the data in the corresponding y, the gray box represents the 1st and 3rd percentiles, the white dot the median, and the whiskers the standard deviation.

### k-means distribution and cluster analysis of stiffness population

The whole procedure was performed in MATLAB. First, the different conditions were merged into an array and plotted to visualize the number of populations. Based on the observation, 4 populations were selected (3 and 6 were also tested, with similar outcomes). Then, k-means was applied to the merged data to get the centroids of the populations and then, these centroids were used to assign a population to each value in the individual condition and used to calculate the weight (%) of each population. The result percentages were used for calculating the Euclidean distances and the Ward algorithm was selected for the linkage, which is represented as a dendrogram. The definition of the number of clusters was based on the inconsistent coefficient.

## Author contributions

Conceptualization: H.R. and M.C.C.; cell line and plasmid generation: M.K.P., N.T. and H.Z.; sample preparation: H.R. and M.K.P.; immunofluorescence and imaging: H.R.; atomic force microscopy (AFM) sampling, measurements and analysis: A.A.; AFM scientific support: A.A and C.D.; RNA/ChIP-seq analysis: P.P.; RT-qPCR: M.A.; Formal analysis: V.B. and H.R; Resources: B.L, R.W.S. and M.C.C.; Funding acquisition: R.W.S. and M.C.C.; Visualization: H.R.; Writing - Original draft: H.R.; Writing - Review and Editing: H.R., A.A, M.A. and M.C.C.

## Material transfer agreements

All data needed to evaluate the conclusions in the manuscript are present in the manuscript and/or the supplementary materials.

All data, included the self-written macros, can be found in the TU-datalib repository (https://tudatalib.ulb.tu-darmstadt.de/handle/tudatalib/4381).

## Supplementary Materials

**Extended Figure S1.**
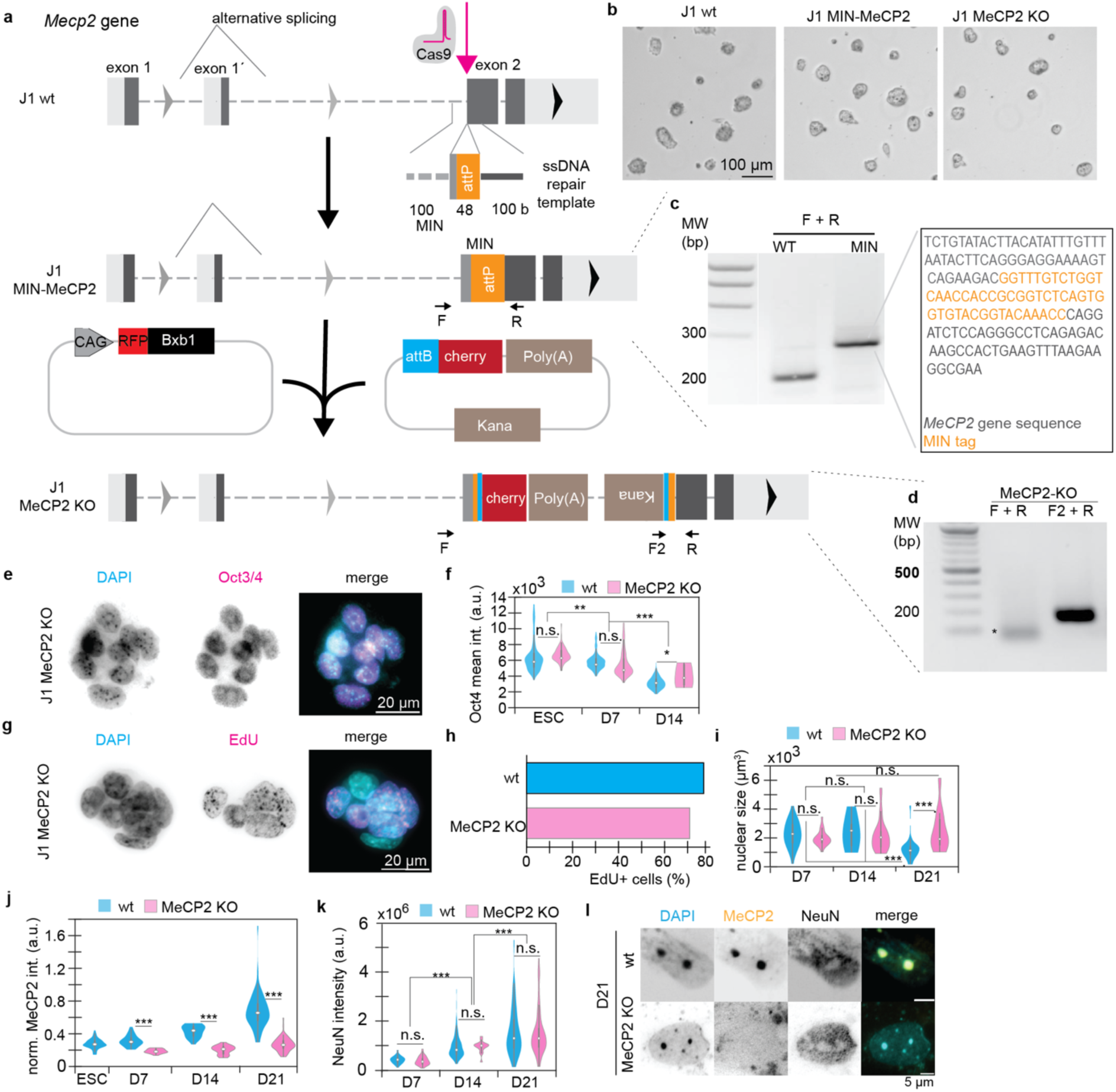
Generation and characterization of *Mecp2* knockout (MeCP2 KO) stem cells and neural differentiation by LIF deprivation. a. Scheme of MIN strategy used for generating the knockout. First, a MIN tag was inserted in the exon 2 of MeCP2 (to knockout both isoforms). Then, transfection with a plasmid containing the recombinase Bxb1 and attB-mCherry-polyA led to the introduction on the plasmid, therefore adding a stop codon in the exon 2. b. Exemplary images of the colony formation of J1, J1 MIN-MeCP2 and J1 MeCP2 KO cells in differential interference contrast images. c. Confirmation of the MIN insertion by PCR screening using the plasmids F-R (Table S2). The expected lengths were 200 bp for the wt and 300 bp. d. Confirmation of the stop codon insertion by double PCR. The expected sizes were ∼5 kb for the F+R and 200 bp for the F2+R. * represent the primed oligos, as the conditions did not allow a complete polymerization of the 5 kb. e-f. Characterization of pluripotency in MeCP2 KO embryonic stem cells by Oct4 immunostaining, showing representative images (e) and the quantification of the intensity of Oct4 normalized to the DNA (DAPI intensity). g-h. Characterization of the replication ability of MeCP2 KO cells by EdU incorporation, showing representative images (g) and the quantification of the EdU positive cells was compared to previous published data in the J1 wt cells (*83*). i. Calculation of the nucleus volume in the DNA staining (DAPI) by 3D confocal microscopy in wt and MeCP2 KO cells. j-l. Characterization of MeCP2 and NeuN levels in the differentiation to neurons by immunostaining. Quantification of MeCP2 levels normalized to DAPI (j) and NeuN levels (k), as well as representative images of the neuronal stage (l). n.s. non-significant; *: p < 0.05; **: p <0.001; ***: p < 0.0001

**Extended Figure S2.**
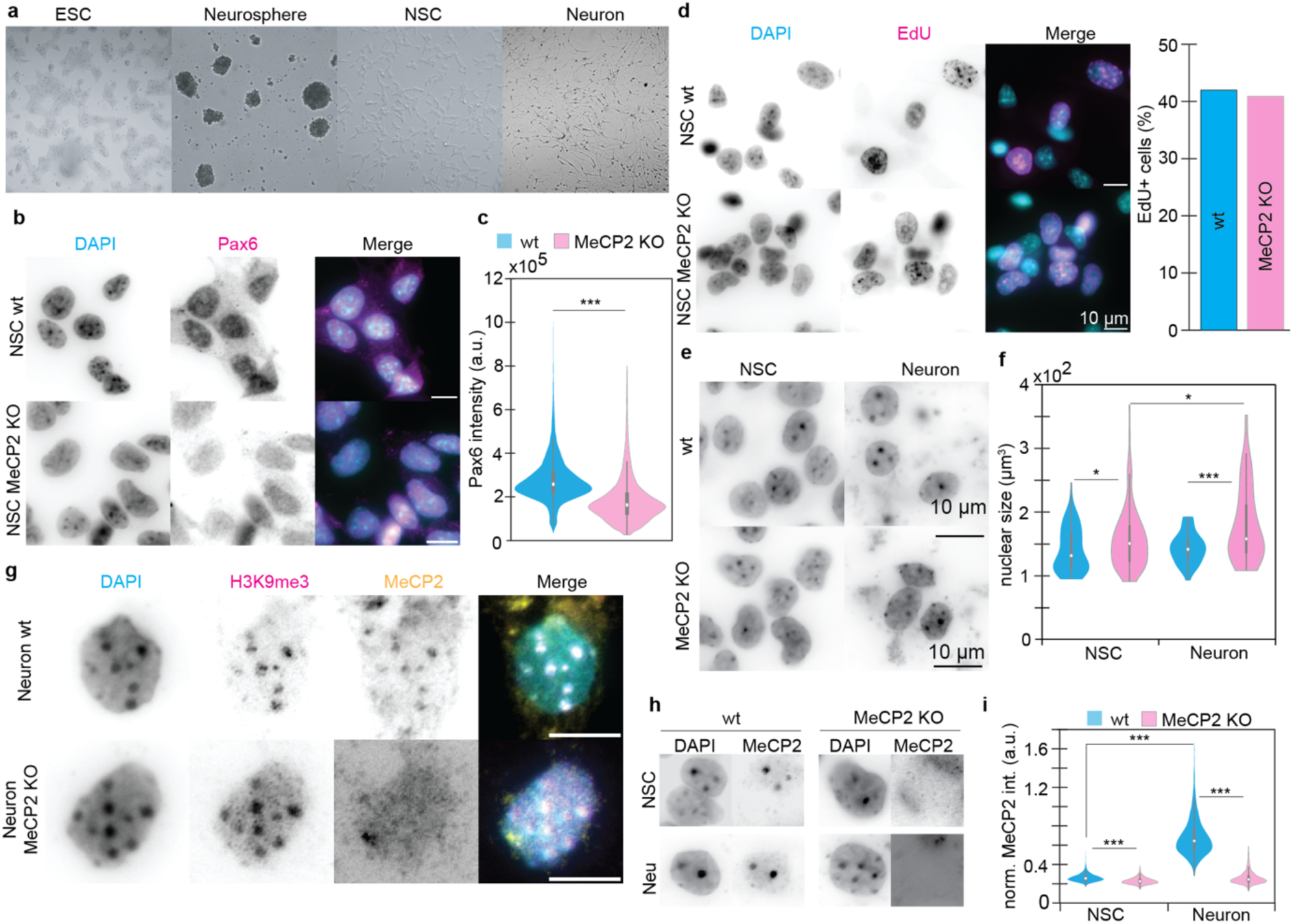
Establishment of regenerating neural stem cells (NSC) from wt and MeCP2 KO ESC and characterization of the differentiation to neurons. a. Overview of the morphology of the cells during the establishment and differentiation. b. Representative images of the NSC marker Pax6 of wt and MeCP2 KO cells. c. Quantification of the levels of Pax6 by high-throughput microscopy. d. Characterization of the self-renewal ability by EdU incorporation and quantification of EdU positive cells in wt and MeCP2 KO cells. e. Overview of the changes in the nucleus in differentiation of NSC to neurons by DNA staining with DAPI. f. Quantification of the nuclear size of NSC and neurons in wt and MeCP2 KO cells by 3D confocal microscopy. g. Confirmation of the MeCP2 KO by immunostaining with MeCP2 together with the heterochromatin marker Histone 3 Lys-9 trimethylation (H3K9me3).

**Extended table S1.** Cell lines used and their characteristics.

| Name | Species | Type | Genotype | Reference |
| --- | --- | --- | --- | --- |
| C2C12 | *Mus musculus* | myoblast | wild type | (92) |
| J1 wt | *Mus musculus* | embryonic stem cell | wild type | (93) |
| J1 MIN-MeCP2 | *Mus musculus* | embryonic stem cell | wild type | this study |
| J1 MeCP2 KO | *Mus musculus* | embryonic stem cell | wild type | this study |
| J1 NSC wt | *Mus musculus* | neural stem cell | wild type | this study |
| J1 NSC MeCP2 KO | *Mus musculus* | neural stem cell | wild type | this study |

**Extended table S2.**
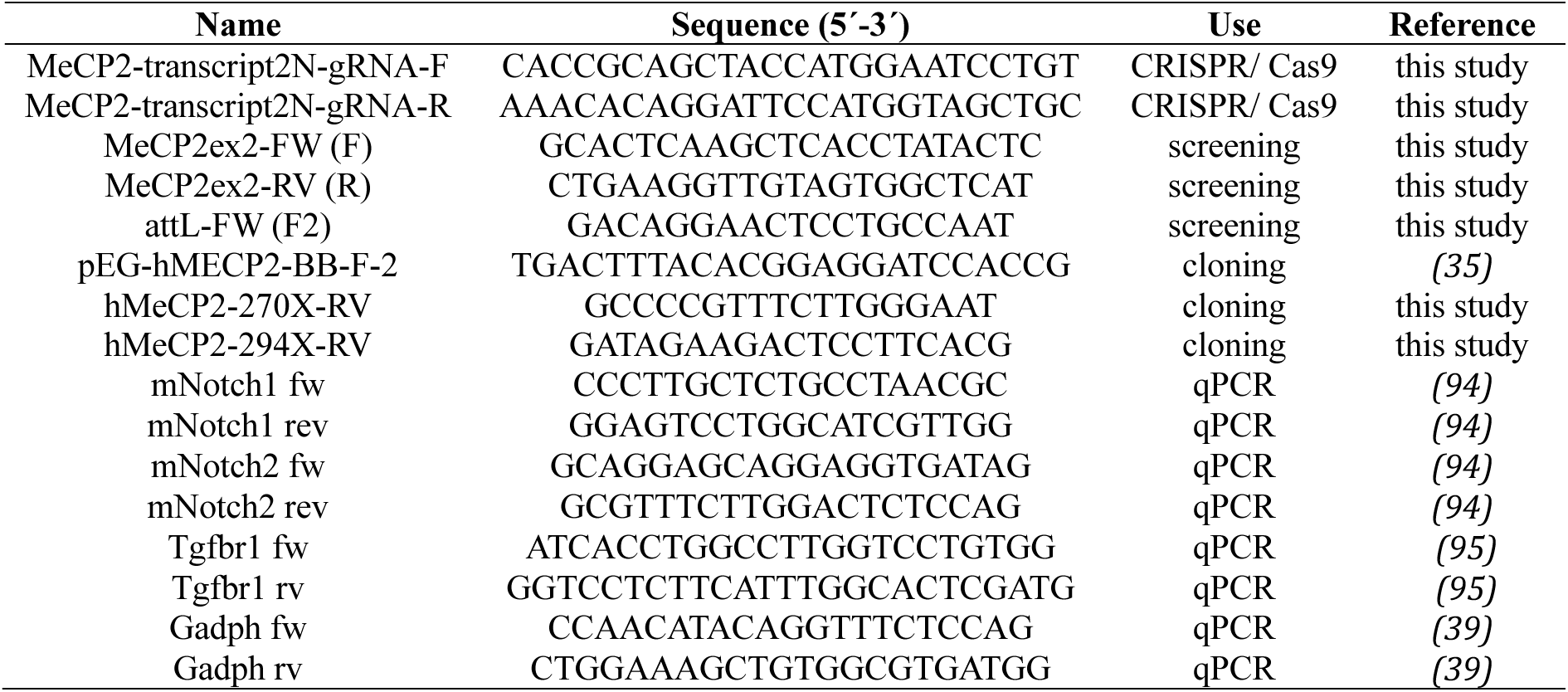
Oligos used in this work.

**Extended table S3.**
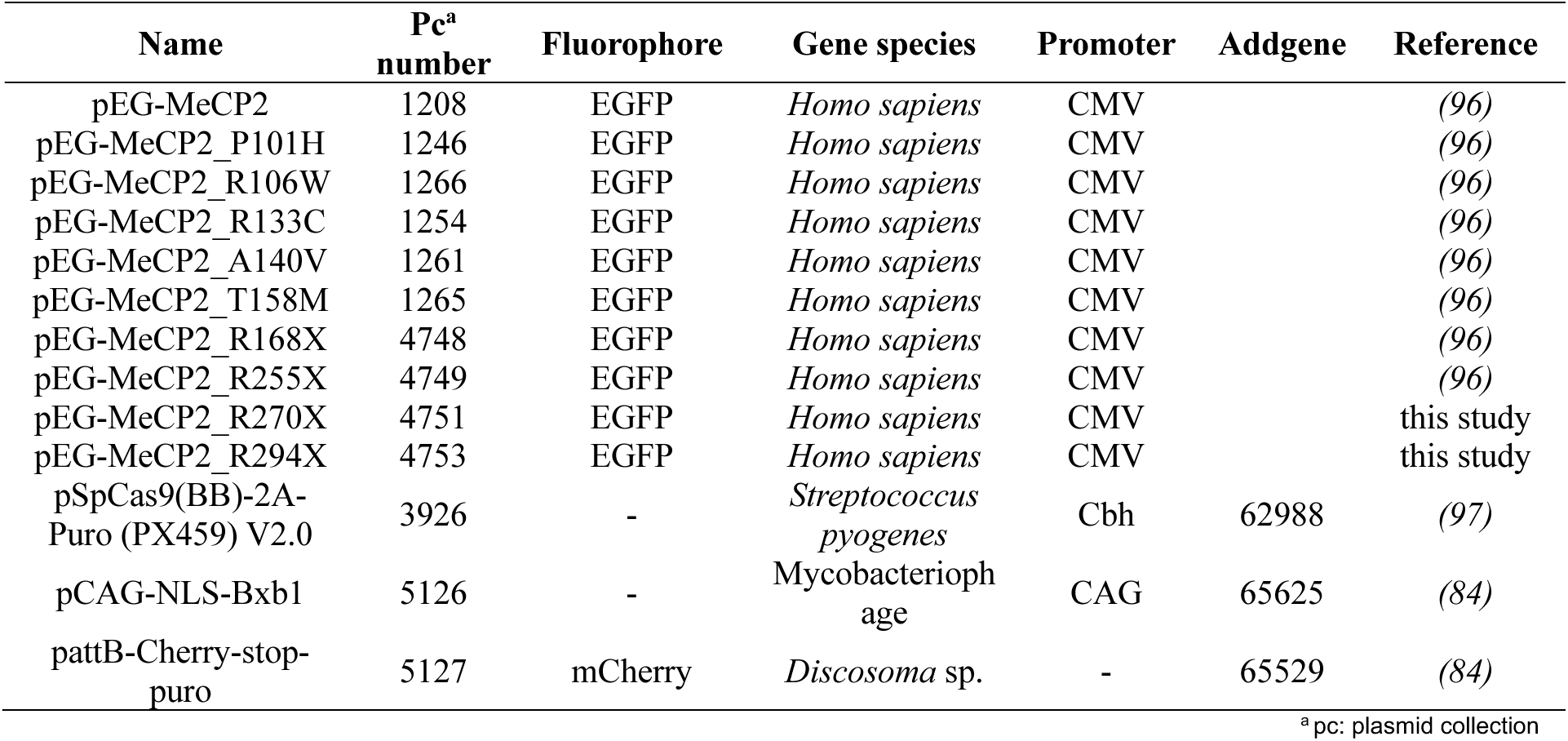
Plasmid used in this work.

| Name | Pc<sup>a</sup><br>number | Fluorophore | Gene species | Promoter | Addgene | Reference |
| --- | --- | --- | --- | --- | --- | --- |
| pEG-MeCP2 | 1208 | EGFP | *Homo sapiens* | CMV |  | (96) |
| pEG-MeCP2_P101H | 1246 | EGFP | *Homo sapiens* | CMV |  | (96) |
| pEG-MeCP2_R106W | 1266 | EGFP | *Homo sapiens* | CMV |  | (96) |
| pEG-MeCP2_R133C | 1254 | EGFP | *Homo sapiens* | CMV |  | (96) |
| pEG-MeCP2_A140V | 1261 | EGFP | *Homo sapiens* | CMV |  | (96) |
| pEG-MeCP2_T158M | 1265 | EGFP | *Homo sapiens* | CMV |  | (96) |
| pEG-MeCP2_R168X | 4748 | EGFP | *Homo sapiens* | CMV |  | (96) |
| pEG-MeCP2_R255X | 4749 | EGFP | *Homo sapiens* | CMV |  | (96) |
| pEG-MeCP2_R270X | 4751 | EGFP | *Homo sapiens* | CMV |  | this study |
| pEG-MeCP2_R294X | 4753 | EGFP | *Homo sapiens* | CMV |  | this study |
| pSpCas9(BB)-2A-Puro (PX459) V2.0 | 3926 | - | *Streptococcus pyogenes* | Cbh | 62988 | (97) |
| pCAG-NLS-Bxb1 | 5126 | - | *Mycobacteriophage* | CAG | 65625 | (84) |
| pattB-Cherry-stop-puro | 5127 | mCherry | *Discosoma* sp. | - | 65529 | (84) |

<sup>a</sup> pc: plasmid collection**Extended table S4.** Antibodies used in this work.

**Extended table S4.** Antibodies used in this work.

| Reactivity | Host | Fixation (IF<sup>a</sup>) | Dilution | Application | Catalog / Clone | Company / Reference |
| --- | --- | --- | --- | --- | --- | --- |
| anti-MeCP2 | rabbit | methanol | 1:250/<br>1:500 | IF<sup>a</sup> / WB<sup>b</sup> | - | (98) |
| anti-MeCP2 | mouse | formaldehyde | undiluted | IF<sup>a</sup> | 4B4 | (98) |
| anti-MeCP2 | rat | formaldehyde /<br>methanol | undiluted | IF<sup>a</sup> | 4H7 | (98) |
| anti-MyoD | mouse | methanol | 1:60 | IF<sup>a</sup> | MoAb5_8a | Pharmigen |
| anti-NeuN | rabbit | formaldehyde | 1:500 | IF<sup>a</sup> | EPR12763 | Abcam |
| anti-Oct3/4 | mouse | formaldehyde | 1:500 | IF<sup>a</sup> | Clone 50/Oct-3<br>(RUO) | BD<br>Biosciences |
| anti-Histone<br>H3 K9<br>trimethyl | rabbit | formaldehyde | 1:200 | IF<sup>a</sup> | 2AG-6F12-H4 | Active Motif |
| anti-Pax6 | rabbit | formaldehyde | 1:100 | IF<sup>a</sup> | 901301 | Biolegend |
| anti-rabbit IgG<br>Alexa Fluor<br>647 | goat | formaldehyde /<br>methanol | 1:400 | IF<sup>a</sup> | 2160406 | Molecular<br>Probes |
| anti-rabbit IgG<br>Cy5 | donkey | formaldehyde /<br>methanol | 1:500 | IF<sup>a</sup> | 711-175-152 | Jackson<br>Immuno<br>Research |
| anti-mouse<br>IgG Cy5 | donkey | formaldehyde /<br>methanol | 1:400 | IF<sup>a</sup> | 711-175-150 | Jackson<br>Immuno<br>Research |
| anti-mouse<br>IgG Alexa<br>Fluor 488 | goat | formaldehyde /<br>methanol | 1:800 | IF<sup>a</sup> | 2120125 | Invitrogen |
| anti-rat IgG<br>Alexa Fluor<br>488 | donkey | formaldehyde /<br>methanol | 1:800 | IF<sup>a</sup> | A_11006 | Invitrogen |

**Extended table S5.** Microscopy and cell sorting devices and their characteristics.

| Microscope / Company | Laser / Lamps | Filters (ex<sup>a</sup> & em<sup>b</sup> [nm]) | Objective / Lenses | Detection system | Application |
| --- | --- | --- | --- | --- | --- |
| S3e Cell Sorter / BioRad laboratories | 488 nm | GFP (em: 525±30) |  | FSC<sup>c</sup> with PMT<sup>d</sup><br>SSC<sup>c</sup> with PMT<sup>d</sup><br>Fluorescence detectors with PMTs<sup>d</sup> | Sort cells according to GFP expression |
| Nikon Eclipse TiE2 / Nikon | Spectra X Led 395±25 (295 mW) 470±24 nm (196 mW) 540±30 nm (231 mW) | em.: Quadbandpass (432±25, 515±25, 595±25, 730±70 nm) | 20X air Splan Fluor LWD DIC (0.7 NA<sup>f</sup>, 2.3 mm WD<sup>g</sup>)<br>40X air Plan Apo λ DIC (0.95 NA<sup>f</sup>, 230 μm WD<sup>g</sup>) | Nikon Qi2 751600 16.25 MPx | Widefield High throughput microscopy |
| Leica TSC SPE-II | Multicolor solid-state laser module RYBC 405 nm (25mW) 488 nm (10 mW) 635 nm (18mW) | em.: DAPI: 410-600 GFP: 493-540 Cy5: 640-783 | ACS APO 63X (1.3 NA<sup>f</sup>, 0.16 WD<sup>g</sup>) | Trans-PMT<sup>d</sup> | 3D confocal |

<sup>a</sup> ex: excitation; <sup>b</sup> em: emission; <sup>c</sup> FSC: forward scatter; <sup>d</sup> PTM: photomultiplier tube(s); <sup>e</sup> SSC: side scatter; <sup>f</sup> NA: numerical aperture; <sup>g</sup> WD: working distance

## References and Citations

1. P. M. Gilbert, K. L. Havenstrite, K. E. G. Magnusson, A. Sacco, N. A. Leonardi, P. Kraft, N. K. Nguyen, S. Thrun, M. P. Lutolf, H. M. Blau, Substrate elasticity regulates skeletal muscle stem cell self-renewal in culture. Science 329, 1078–1081 (2010).

2. V. Venturini, F. Pezzano, F. Català Castro, H.-M. Häkkinen, S. Jiménez-Delgado, M. Colomer-Rosell, M. Marro, Q. Tolosa-Ramon, S. Paz-López, M. A. Valverde, J. Weghuber, P. Loza-Alvarez, M. Krieg, S. Wieser, V. Ruprecht, The nucleus measures shape changes for cellular proprioception to control dynamic cell behavior. Science 370 (2020).

3. N. D. Evans, C. Minelli, E. Gentleman, V. LaPointe, S. N. Patankar, M. Kallivretaki, X. Chen, C. J. Roberts, M. M. Stevens, Substrate stiffness affects early differentiation events in embryonic stem cells. Eur. Cell. Mater. 18, 1–13; discussion 13 (2009).

4. A. J. Engler, S. Sen, H. L. Sweeney, D. E. Discher, Matrix elasticity directs stem cell lineage specification. Cell 126, 677–689 (2006).

5. S. Ali, I. B. Wall, C. Mason, A. E. Pelling, F. S. Veraitch, The effect of Young’s modulus on the neuronal differentiation of mouse embryonic stem cells. Acta Biomater. 25, 253–267 (2015).

6. A. Kumar, J. K. Placone, A. J. Engler, Understanding the extracellular forces that determine cell fate and maintenance. Development 144, 4261–4270 (2017).

7. X. Zhang, S. Zhang, T. Wang, How the mechanical microenvironment of stem cell growth affects their differentiation: a review. Stem Cell Res. Ther. 13, 415 (2022).

8. J. D. Pajerowski, K. N. Dahl, F. L. Zhong, P. J. Sammak, D. E. Discher, Physical plasticity of the nucleus in stem cell differentiation. Proc Natl Acad Sci USA 104, 15619–15624 (2007).

9. C. M. Hall, E. Moeendarbary, G. K. Sheridan, Mechanobiology of the brain in ageing and Alzheimer’s disease. Eur. J. Neurosci. 53, 3851–3878 (2021).

10. L. V. Hiscox, C. L. Johnson, M. D. J. McGarry, H. Marshall, C. W. Ritchie, E. J. R. van Beek, N. Roberts, J. M. Starr, Mechanical property alterations across the cerebral cortex due to Alzheimer’s disease. Brain Commun. 2, fcz049 (2020).

11. J. Swift, I. L. Ivanovska, A. Buxboim, T. Harada, P. C. D. P. Dingal, J. Pinter, J. D. Pajerowski, K. R. Spinler, J.-W. Shin, M. Tewari, F. Rehfeldt, D. W. Speicher, D. E. Discher, Nuclear lamin-A scales with tissue stiffness and enhances matrix-directed differentiation. Science 341, 1240104 (2013).

12. A. Procès, M. Luciano, Y. Kalukula, L. Ris, S. Gabriele, Multiscale mechanobiology in brain physiology and diseases. Front. Cell Dev. Biol. 10, 823857 (2022).

13. E. K. F. Yim, M. P. Sheetz, Force-dependent cell signaling in stem cell differentiation. Stem Cell Res. Ther. 3, 41 (2012).

14. W. J. Tyler, The mechanobiology of brain function. Nat. Rev. Neurosci. 13, 867–878 (2012).

15. M. Lam, F. Calvo, Regulation of mechanotransduction: Emerging roles for septins. Cytoskeleton (Hoboken) 76, 115–122 (2019).

16. C. S. Janota, F. J. Calero-Cuenca, E. R. Gomes, The role of the cell nucleus in mechanotransduction. Curr. Opin. Cell Biol. 63, 204–211 (2020).

17. D. Amiad Pavlov, C. P. Unnikannan, D. Lorber, G. Bajpai, T. Olender, E. Stoops, A. Reuveny, S. Safran, T. Volk, The LINC complex inhibits excessive chromatin repression. BioRxiv (2022).

18. A. Piscioneri, S. Morelli, T. Ritacco, M. Giocondo, R. Peñaloza, E. Drioli, L. De Bartolo, Topographical cues of PLGA membranes modulate the behavior of hMSCs, myoblasts and neuronal cells. Colloids Surf. B Biointerfaces 222, 113070 (2023).

19. M. M. Nava, Y. A. Miroshnikova, L. C. Biggs, D. B. Whitefield, F. Metge, J. Boucas, H. Vihinen, E. Jokitalo, X. Li, J. M. García Arcos, B. Hoffmann, R. Merkel, C. M. Niessen, K. N. Dahl, S. A. Wickström, Heterochromatin-Driven Nuclear Softening Protects the Genome against Mechanical Stress-Induced Damage. Cell 181, 800–817.e22 (2020).

20. K. N. Dahl, A. J. S. Ribeiro, J. Lammerding, Nuclear shape, mechanics, and mechanotransduction. Circ. Res. 102, 1307–1318 (2008).

21. K. N. Dahl, G. W. G. Luxton, A special topic on nuclear mechanobiology. Cell. Mol. Bioeng. 9, 203–206 (2016).

22. D. Amiad-Pavlov, D. Lorber, G. Bajpai, A. Reuveny, F. Roncato, R. Alon, S. Safran, T. Volk, Live imaging of chromatin distribution reveals novel principles of nuclear architecture and chromatin compartmentalization. Sci. Adv. 7 (2021).

23. A. Vahabikashi, S. Sivagurunathan, F. A. S. Nicdao, Y. L. Han, C. Y. Park, M. Kittisopikul, X. Wong, J. R. Tran, G. G. Gundersen, K. L. Reddy, G. W. G. Luxton, M. Guo, J. J. Fredberg, Y. Zheng, S. A. Adam, R. D. Goldman, Nuclear lamin isoforms differentially contribute to LINC complex-dependent nucleocytoskeletal coupling and whole-cell mechanics. Proc Natl Acad Sci USA 119, e2121816119 (2022).

24. A. R. Killaars, C. J. Walker, K. S. Anseth, Nuclear mechanosensing controls MSC osteogenic potential through HDAC epigenetic remodeling. Proc Natl Acad Sci USA 117, 21258–21266 (2020).

25. A. R. Killaars, J. C. Grim, C. J. Walker, E. A. Hushka, T. E. Brown, K. S. Anseth, Extended exposure to stiff microenvironments leads to persistent chromatin remodeling in human mesenchymal stem cells. Adv Sci (Weinh) 6, 1801483 (2019).

26. W. J. Hadden, J. L. Young, A. W. Holle, M. L. McFetridge, D. Y. Kim, P. Wijesinghe, H. Taylor-Weiner, J. H. Wen, A. R. Lee, K. Bieback, B.-N. Vo, D. D. Sampson, B. F. Kennedy, J. P. Spatz, A. J. Engler, Y. S. Choi, Stem cell migration and mechanotransduction on linear stiffness gradient hydrogels. Proc Natl Acad Sci USA 114, 5647–5652 (2017).

27. T. L. Downing, J. Soto, C. Morez, T. Houssin, A. Fritz, F. Yuan, J. Chu, S. Patel, D. V. Schaffer, S. Li, Biophysical regulation of epigenetic state and cell reprogramming. Nat. Mater. 12, 1154–1162 (2013).

28. E. Makhija, D. S. Jokhun, G. V. Shivashankar, Nuclear deformability and telomere dynamics are regulated by cell geometric constraints. Proc Natl Acad Sci USA 113, E32–40 (2016).

29. Y. Wang, M. Nagarajan, C. Uhler, G. V. Shivashankar, Orientation and repositioning of chromosomes correlate with cell geometry-dependent gene expression. Mol. Biol. Cell 28, 1997–2009 (2017).

30. F. Alisafaei, D. S. Jokhun, G. V. Shivashankar, V. B. Shenoy, Regulation of nuclear architecture, mechanics, and nucleocytoplasmic shuttling of epigenetic factors by cell geometric constraints. Proc Natl Acad Sci USA 116, 13200–13209 (2019).

31. I. Solovei, M. Kreysing, C. Lanctôt, S. Kösem, L. Peichl, T. Cremer, J. Guck, B. Joffe, Nuclear architecture of rod photoreceptor cells adapts to vision in mammalian evolution. Cell 137, 356–368 (2009).

32. R. Terranova, S. Sauer, M. Merkenschlager, A. G. Fisher, The reorganisation of constitutive heterochromatin in differentiating muscle requires HDAC activity. Exp. Cell Res. 310, 344–356 (2005).

33. A. Brero, H. P. Easwaran, D. Nowak, I. Grunewald, T. Cremer, H. Leonhardt, M. C. Cardoso, Methyl CpG-binding proteins induce large-scale chromatin reorganization during terminal differentiation. J. Cell Biol. 169, 733–743 (2005).

34. B. Bertulat, M. L. De Bonis, F. Della Ragione, A. Lehmkuhl, M. Milden, C. Storm, K. L. Jost, S. Scala, B. Hendrich, M. D’Esposito, M. C. Cardoso, MeCP2 dependent heterochromatin reorganization during neural differentiation of a novel Mecp2-deficient embryonic stem cell reporter line. PLoS ONE 7, e47848 (2012).

35. H. Zhang, H. Romero, A. Schmidt, K. Gagova, W. Qin, B. Bertulat, A. Lehmkuhl, M. Milden, M. Eck, T. Meckel, H. Leonhardt, M. C. Cardoso, MeCP2-induced heterochromatin organization is driven by oligomerization-based liquid-liquid phase separation and restricted by DNA methylation. Nucleus 13, 1–34 (2022).

36. A. Schmidt, H. Zhang, M. C. Cardoso, MeCP2 and chromatin compartmentalization. Cells 9 (2020).

37. R. E. Amir, I. B. Van den Veyver, M. Wan, C. Q. Tran, U. Francke, H. Y. Zoghbi, Rett syndrome is caused by mutations in X-linked MECP2, encoding methyl-CpG-binding protein 2. Nat. Genet. 23, 185–188 (1999).

38. J. Guy, B. Hendrich, M. Holmes, J. E. Martin, A. Bird, A mouse Mecp2-null mutation causes neurological symptoms that mimic Rett syndrome. Nat. Genet. 27, 322–326 (2001).

39. P. J. Skene, R. S. Illingworth, S. Webb, A. R. W. Kerr, K. D. James, D. J. Turner, R. Andrews, A. P. Bird, Neuronal MeCP2 is expressed at near histone-octamer levels and globally alters the chromatin state. Mol. Cell 37, 457–468 (2010).

40. C. Song, Y. Feodorova, J. Guy, L. Peichl, K. L. Jost, H. Kimura, M. C. Cardoso, A. Bird, H. Leonhardt, B. Joffe, I. Solovei, DNA methylation reader MECP2: cell type- and differentiation stage-specific protein distribution. Epigenetics Chromatin 7, 17 (2014).

41. G. Bajpai, D. Amiad Pavlov, D. Lorber, T. Volk, S. Safran, Mesoscale phase separation of chromatin in the nucleus. eLife 10 (2021).

42. H. Romero, A. Schmidt, C. M. Cardoso, Protein Level Quantification Across Fluorescence-based Platforms. Bio Protoc 13, e4834 (2023).

43. A. Pillarisetti, J. P. Desai, H. Ladjal, A. Schiffmacher, A. Ferreira, C. L. Keefer, Mechanical phenotyping of mouse embryonic stem cells: increase in stiffness with differentiation. Cell. Reprogram. 13, 371–380 (2011).

44. M. Yazdani, R. Deogracias, J. Guy, R. A. Poot, A. Bird, Y.-A. Barde, Disease modeling using embryonic stem cells: MeCP2 regulates nuclear size and RNA synthesis in neurons. Stem Cells 30, 2128–2139 (2012).

45. C. Philippe, L. Villard, N. De Roux, M. Raynaud, J. P. Bonnefond, L. Pasquier, G. Lesca, J. Mancini, P. Jonveaux, A. Moncla, J. Chelly, T. Bienvenu, Spectrum and distribution of MECP2 mutations in 424 Rett syndrome patients: a molecular update. Eur. J. Med. Genet. 49, 9–18 (2006).

46. R. Krishnaraj, G. Ho, J. Christodoulou, RettBASE: Rett syndrome database update. Hum. Mutat. 38, 922–931 (2017).

47. T. Fukuda, Y. Yamashita, S. Nagamitsu, K. Miyamoto, J.-J. Jin, I. Ohmori, Y. Ohtsuka, K. Kuwajima, S. Endo, T. Iwai, H. Yamagata, Y. Tabara, T. Miki, T. Matsuishi, I. Kondo, Methyl-CpG binding protein 2 gene (MECP2) variations in Japanese patients with Rett syndrome: pathological mutations and polymorphisms. Brain Dev. 27, 211–217 (2005).

48. N. Agarwal, A. Becker, K. L. Jost, S. Haase, B. K. Thakur, A. Brero, T. Hardt, S. Kudo, H. Leonhardt, M. C. Cardoso, MeCP2 Rett mutations affect large scale chromatin organization. Hum. Mol. Genet. 20, 4187–4195 (2011).

49. K. P. McCreery, X. Xu, A. K. Scott, A. K. Fajrial, S. Calve, X. Ding, C. P. Neu, Nuclear Stiffness Decreases with Disruption of the Extracellular Matrix in Living Tissues. Small 17, e2006699 (2021).

50. K. N. Dahl, S. M. Kahn, K. L. Wilson, D. E. Discher, The nuclear envelope lamina network has elasticity and a compressibility limit suggestive of a molecular shock absorber. J. Cell Sci. 117, 4779–4786 (2004).

51. J. Lammerding, P. C. Schulze, T. Takahashi, S. Kozlov, T. Sullivan, R. D. Kamm, C. L. Stewart, R. T. Lee, Lamin A/C deficiency causes defective nuclear mechanics and mechanotransduction. J. Clin. Invest. 113, 370–378 (2004).

52. T. Fischer, A. Hayn, C. T. Mierke, Effect of nuclear stiffness on cell mechanics and migration of human breast cancer cells. Front. Cell Dev. Biol. 8, 393 (2020).

53. A. Vahabikashi, S. A. Adam, O. Medalia, R. D. Goldman, Nuclear lamins: Structure and function in mechanobiology. APL Bioengineering 6, 011503 (2022).

54. A. D. Stephens, E. J. Banigan, S. A. Adam, R. D. Goldman, J. F. Marko, Chromatin and lamin A determine two different mechanical response regimes of the cell nucleus. Mol. Biol. Cell 28, 1984–1996 (2017).

55. A. Elosegui-Artola, I. Andreu, A. E. M. Beedle, A. Lezamiz, M. Uroz, A. J. Kosmalska, R. Oria, J. Z. Kechagia, P. Rico-Lastres, A.-L. Le Roux, C. M. Shanahan, X. Trepat, D. Navajas, S. Garcia-Manyes, P. Roca-Cusachs, Force Triggers YAP Nuclear Entry by Regulating Transport across Nuclear Pores. Cell 171, 1397–1410.e14 (2017).

56. I. Andreu, I. Granero-Moya, S. Garcia-Manyes, P. Roca-Cusachs, Understanding the role of mechanics in nucleocytoplasmic transport. APL Bioengineering 6, 020901 (2022).

57. E. C. Jacobson, J. K. Perry, D. S. Long, A. L. Olins, D. E. Olins, B. E. Wright, M. H. Vickers, J. M. O’Sullivan, Migration through a small pore disrupts inactive chromatin organization in neutrophil-like cells. BMC Biol. 16, 142 (2018).

58. N. M. Ramdas, G. V. Shivashankar, Cytoskeletal control of nuclear morphology and chromatin organization. J. Mol. Biol. 427, 695–706 (2015).

59. N. Ballas, D. T. Lioy, C. Grunseich, G. Mandel, Non-cell autonomous influence of MeCP2-deficient glia on neuronal dendritic morphology. Nat. Neurosci. 12, 311–317 (2009).

60. N. Kishi, J. D. Macklis, MECP2 is progressively expressed in post-migratory neurons and is involved in neuronal maturation rather than cell fate decisions. Mol. Cell. Neurosci. 27, 306–321 (2004).

61. M. V. Johnston, M. E. Blue, S. Naidu, Rett syndrome and neuronal development. J. Child Neurol. 20, 759–763 (2005).

62. S. C. Sweat, C. E. J. Cheetham, Deficits in olfactory system neurogenesis in neurodevelopmental disorders. Genesis 62, e23590 (2024).

63. M. K. Singleton, M. L. Gonzales, K. N. Leung, D. H. Yasui, D. I. Schroeder, K. Dunaway, J. M. LaSalle, MeCP2 is required for global heterochromatic and nucleolar changes during activity-dependent neuronal maturation. Neurobiol. Dis. 43, 190–200 (2011).

64. C. Engel-Pizcueta, C. Pujades, Interplay between notch and YAP/TAZ pathways in the regulation of cell fate during embryo development. Front. Cell Dev. Biol. 9, 711531 (2021).

65. A. Engler, C. Rolando, C. Giachino, I. Saotome, A. Erni, C. Brien, R. Zhang, U. Zimber-Strobl, F. Radtke, S. Artavanis-Tsakonas, A. Louvi, V. Taylor, Notch2 Signaling Maintains NSC Quiescence in the Murine Ventricular-Subventricular Zone. Cell Rep. 22, 992–1002 (2018).

66. D. J. Solecki, X. L. Liu, T. Tomoda, Y. Fang, M. E. Hatten, Activated Notch2 signaling inhibits differentiation of cerebellar granule neuron precursors by maintaining proliferation. Neuron 31, 557–568 (2001).

67. Y. He, H. Zhang, A. Yung, S. A. Villeda, P. A. Jaeger, O. Olayiwola, N. Fainberg, T. Wyss-Coray, ALK5-dependent TGF-β signaling is a major determinant of late-stage adult neurogenesis. Nat. Neurosci. 17, 943–952 (2014).

68. A. R. Gomes, T. G. Fernandes, S. H. Vaz, T. P. Silva, E. P. Bekman, S. Xapelli, S. Duarte, M. Ghazvini, J. Gribnau, A. R. Muotri, C. A. Trujillo, A. M. Sebastião, J. M. S. Cabral, M. M. Diogo, Modeling Rett Syndrome With Human Patient-Specific Forebrain Organoids. Front. Cell Dev. Biol. 8, 610427 (2020).

69. R. D. Smrt, J. Eaves-Egenes, B. Z. Barkho, N. J. Santistevan, C. Zhao, J. B. Aimone, F. H. Gage, X. Zhao, Mecp2 deficiency leads to delayed maturation and altered gene expression in hippocampal neurons. Neurobiol. Dis. 27, 77–89 (2007).

70. K.-Y. Kim, E. Hysolli, I.-H. Park, Neuronal maturation defect in induced pluripotent stem cells from patients with Rett syndrome. Proc Natl Acad Sci USA 108, 14169–14174 (2011).

71. M. Klüppel, J. L. Wrana, Turning it up a Notch: cross-talk between TGF beta and Notch signaling. Bioessays 27, 115–118 (2005).

72. J. Zavadil, L. Cermak, N. Soto-Nieves, E. P. Böttinger, Integration of TGF-beta/Smad and Jagged1/Notch signalling in epithelial-to-mesenchymal transition. EMBO J. 23, 1155–1165 (2004).

73. A. Blokzijl, C. Dahlqvist, E. Reissmann, A. Falk, A. Moliner, U. Lendahl, C. F. Ibáñez, Cross-talk between the Notch and TGF-beta signaling pathways mediated by interaction of the Notch intracellular domain with Smad3. J. Cell Biol. 163, 723–728 (2003).

74. J. Su, L. Guo, C. Wu, A mechanoresponsive PINCH-1-Notch2 interaction regulates smooth muscle differentiation of human placental mesenchymal stem cells. Stem Cells 39, 650–668 (2021).

75. T. Babushku, M. Lechner, S. Ehrenberg, U. Rambold, M. Schmidt-Supprian, A. J. Yates, S. Rane, U. Zimber-Strobl, L. J. Strobl, Notch2 controls developmental fate choices between germinal center and marginal zone B cells upon immunization. Nat. Commun. 15, 1960 (2024).

76. M. M. G. Hillege, A. Shi, R. A. Galli, G. Wu, P. Bertolino, W. M. H. Hoogaars, R. T. Jaspers, Lack of Tgfbr1 and Acvr1b synergistically stimulates myofibre hypertrophy and accelerates muscle regeneration. eLife 11 (2022).

77. Y. Gao, K. J. Bayless, Q. Li, TGFBR1 is required for mouse myometrial development. Mol. Endocrinol. 28, 380–394 (2014).

78. V. A. Cuddapah, R. B. Pillai, K. V. Shekar, J. B. Lane, K. J. Motil, S. A. Skinner, D. C. Tarquinio, D. G. Glaze, G. McGwin, W. E. Kaufmann, A. K. Percy, J. L. Neul, M. L. Olsen, Methyl-CpG-binding protein 2 (MECP2) mutation type is associated with disease severity in Rett syndrome. J. Med. Genet. 51, 152–158 (2014).

79. F. S. Pidcock, C. Salorio, G. Bibat, J. Swain, J. Scheller, W. Shore, S. Naidu, Functional outcomes in Rett syndrome. Brain Dev. 38, 76–81 (2016).

80. K. Brown, J. Selfridge, S. Lagger, J. Connelly, D. De Sousa, A. Kerr, S. Webb, J. Guy, C. Merusi, M. V. Koerner, A. Bird, The molecular basis of variable phenotypic severity among common missense mutations causing Rett syndrome. Hum. Mol. Genet. 25, 558–570 (2016).

81. Y. Yang, T. G. Kucukkal, J. Li, E. Alexov, W. Cao, Binding Analysis of Methyl-CpG Binding Domain of MeCP2 and Rett Syndrome Mutations. ACS Chem. Biol. 11, 2706–2715 (2016).

82. E. Ballestar, T. M. Yusufzai, A. P. Wolffe, Effects of Rett syndrome mutations of the methyl-CpG binding domain of the transcriptional repressor MeCP2 on selectivity for association with methylated DNA. Biochemistry 39, 7100–7106 (2000).

83. C. Rausch, P. Weber, P. Prorok, D. Hörl, A. Maiser, A. Lehmkuhl, V. O. Chagin, C. S. Casas-Delucchi, H. Leonhardt, M. C. Cardoso, Developmental differences in genome replication program and origin activation. Nucleic Acids Res. 48, 12751–12777 (2020).

84. C. B. Mulholland, M. Smets, E. Schmidtmann, S. Leidescher, Y. Markaki, M. Hofweber, W. Qin, M. Manzo, E. Kremmer, K. Thanisch, C. Bauer, P. Rombaut, F. Herzog, H. Leonhardt, S. Bultmann, A modular open platform for systematic functional studies under physiological conditions. Nucleic Acids Res. 43, e112 (2015).

85. A. Nabbi, K. Riabowol, Rapid Isolation of Nuclei from Cells In Vitro. Cold Spring Harb. Protoc. 2015, 769–772 (2015).

86. H. J. Butt, M. Jaschke, Calculation of thermal noise in atomic force microscopy. Nanotechnology 6, 1–7 (1995).

87. L. Stühn, A. Fritschen, J. Choy, M. Dehnert, C. Dietz, Nanomechanical sub-surface mapping of living biological cells by force microscopy. Nanoscale 11, 13089–13097 (2019).

88. R. Dougherty, paper presented at 11th AIAA/CEAS Aeroacoustics Conference, 11th AIAA/CEAS Aeroacoustics Conference, 23 May 2005.

89. M. E. Wynne, A. R. Lane, K. S. Singleton, S. A. Zlatic, A. Gokhale, E. Werner, D. Duong, J. Q. Kwong, A. J. Crocker, V. Faundez, Heterogeneous expression of nuclear encoded mitochondrial genes distinguishes inhibitory and excitatory neurons. eNeuro 8 (2021).

90. B. S. Johnson, Y.-T. Zhao, M. Fasolino, J. M. Lamonica, Y. J. Kim, G. Georgakilas, K. H. Wood, D. Bu, Y. Cui, D. Goffin, G. Vahedi, T. H. Kim, Z. Zhou, Biotin tagging of MeCP2 in mice reveals contextual insights into the Rett syndrome transcriptome. Nat. Med. 23, 1203–1214 (2017).

91. B. Kinde, D. Y. Wu, M. E. Greenberg, H. W. Gabel, DNA methylation in the gene body influences MeCP2-mediated gene repression. Proc Natl Acad Sci USA 113, 15114–15119 (2016).

92. D. Yaffe, O. Saxel, Serial passaging and differentiation of myogenic cells isolated from dystrophic mouse muscle. Nature 270, 725–727 (1977).

93. E. Li, T. H. Bestor, R. Jaenisch, Targeted mutation of the DNA methyltransferase gene results in embryonic lethality. Cell 69, 915–926 (1992).

94. N. Trujillo-Paredes, C. Valencia, G. Guerrero-Flores, D.-M. Arzate, J.-M. Baizabal, M. Guerra-Crespo, A. Fuentes-Hernández, I. Zea-Armenta, L. Covarrubias, Regulation of differentiation flux by Notch signalling influences the number of dopaminergic neurons in the adult brain. Biol. Open 5, 336–347 (2016).

95. D. Pohlers, A. Beyer, D. Koczan, T. Wilhelm, H.-J. Thiesen, R. W. Kinne, Constitutive upregulation of the transforming growth factor-beta pathway in rheumatoid arthritis synovial fibroblasts. Arthritis Res. Ther. 9, R59 (2007).

96. S. Kudo, Y. Nomura, M. Segawa, N. Fujita, M. Nakao, C. Schanen, M. Tamura, Heterogeneity in residual function of MeCP2 carrying missense mutations in the methyl CpG binding domain. J. Med. Genet. 40, 487–493 (2003).

97. F. A. Ran, P. D. Hsu, J. Wright, V. Agarwala, D. A. Scott, F. Zhang, Genome engineering using the CRISPR-Cas9 system. Nat. Protoc. 8, 2281–2308 (2013).

98. K. L. Jost, A. Rottach, M. Milden, B. Bertulat, A. Becker, P. Wolf, J. Sandoval, P. Petazzi, D. Huertas, M. Esteller, E. Kremmer, H. Leonhardt, M. C. Cardoso, Generation and characterization of rat and mouse monoclonal antibodies specific for MeCP2 and their use in X-inactivation studies. PLoS ONE 6, e26499 (2011).

